# Springtail-inspired compliant hinge enables terrain-adaptable takeoff in insect-scale robots

**DOI:** 10.64898/2026.02.04.703923

**Authors:** Jacob S. Harrison, Baekgyeom Kim, Hungtang Ko, Adrian A. Smith, Thu Truong, Je-sung Koh, Saad Bhamla

## Abstract

Springtails execute millisecond-scale escape jumps with a single appendage, the furca, on soil, snow, leaf litter, and water. Across 15 taxonomic families (n=552 individuals), relative furca length is bimodal. High-speed video and confocal imaging show that in some long-furca springtails, the resilin-rich manubrium-dens joint behaves as a compliant hinge. It bends during push-off to prolong contact, suppress pitch, and bias takeoff forward, whereas rigid joints drive backward launches with rapid body rotation. We translate this mechanism to a 20-mm, 84-mg jumping robot with an elastic robo-furca hinge. This flexible hinge reduces body rotation by ∼ 90% on flat ground compared to rigid-hinge designs, while maintaining takeoff speed on gravel, springboards, leaves, and pine needles, enabling passive, terrain-adaptable launches for power-limited insect-scale robots without onboard sensing or active control.

## Introduction

In natural settings, jumping animals must navigate uneven and unpredictable substrates to evade predator attacks or reach new resources (*1–5*). Springtails (Collembola; 1-5 mm), among the most abundant arthropods in soil ecosystems, are a global, diverse group of hexapods that use a specialized jumping appendage, the furca (or furcula), to execute ultrafast jumps on soil, sand, mud, snow, rocks, leaves, and even water (Fig. 1A) (*6–11*). To jump, the furca is held beneath the body by retinaculum and then rapidly released to strike the substrate, launching the animal away from threats or toward new habitat (*6, 12, 13*). Because the furca is the sole propulsive appendage, springtails let us isolate how geometry and compliance set launch direction and pitch, in a clade that routinely encounters complex natural substrates (Fig. 1B).

**Figure 1:**
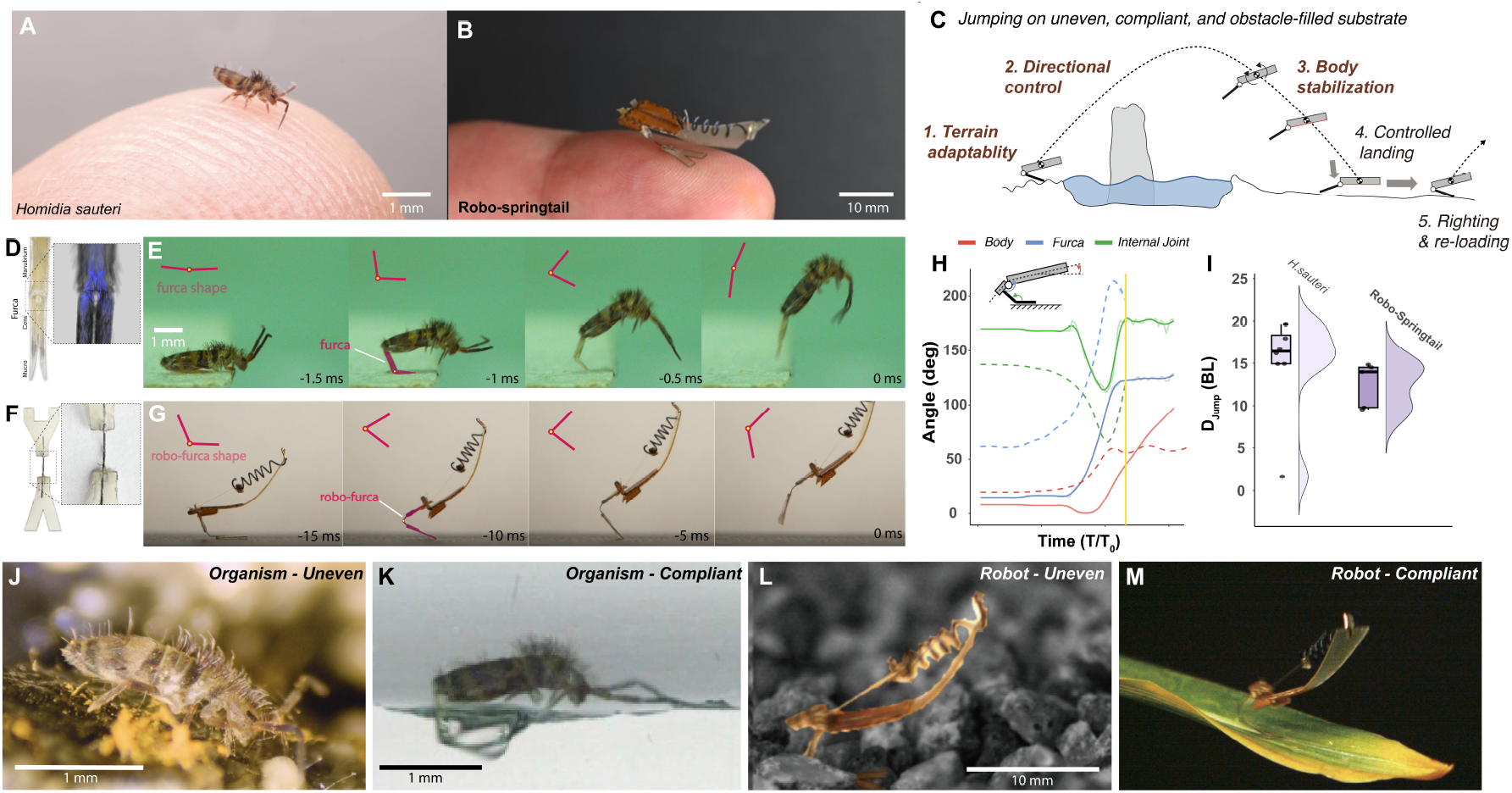
Compliant furcas enable ultrafast jumps on variable substrates in springtails and robots. (A) The springtail *H. sauteri*, a long-compliant focal species, on a fingertip. (B) Long-compliant robo-springtail on a finger. (C) Schematic of the coupled phases required for jumping on uneven, compliant, and obstacle-filled terrain: terrain-adaptive takeoff, directional control, aerial stabilization, controlled landing, and righting/reloading. (D) Furca anatomy (manubrium, dens, and mucro) with an inset confocal image of the manubrium-dens joint; blue fluorescence indicates resilin-rich cuticle. (E) High-speed takeoff sequence of springtail; purple markers indicate furca angle. (F) Robo-furca (jumping appendage) with a flexible joint formed using superelastic nitinol wire. (G) High-speed takeoff sequence of the robo-springtail; purple markers indicate robo-furca angle. (H) Body, furca, and internal furca joint angles during takeoff for *H. sauteri* (solid) and the long-compliant robo-springtail (dashed). (I) Jump distance in body lengths for *H. sauteri* and the long-compliant robotic springtail. (J-M) *H*.*sauteri* and the long-compliant robo-springtail on uneven and deformable natural substrates.

Engineered jumpers face the same constraint. Irregular or deformable ground promotes slip and substrate flexure, which dissipate stored energy during push-off, degrading takeoff speed and stability in both animals and robots (Fig. 1C) (*14, 15*). Larger robotic systems (>100 g and 15 cm) can, in principle, compensate with active control or multiple actuators that adjust body posture and leg motion during takeoff (*16, 17*). At the insect scale, however, tight constraints on size, mass, and power make such active strategies difficult to implement (*18*). Small jumping robots must solve several coupled control challenges — terrain adaptability, directional control, aerial stabilization, landing, and reloading — under millisecond timescales where feedback is limited (Fig. 1C and Table S1). Recent robots have begun to address some of these challenges by using latch designs to regulate takeoff timing on compliant substrates (*19*) or by adding aerodynamic surfaces and mass distributions that improve aerial righting and landing (*11*). Nonetheless, there remains a need for passive mechanical elements that, without additional actuators, can simultaneously improve terrain adaptability and directional control in insect-scale jumping robots.

Here, we ask how furca length and distal compliance shape escape jumps in springtails, and whether a similar elastic hinge can provide terrain-adaptable takeoff at the insect scale. Springtail species differ in the structure and flexibility of their furca. Some have a long furca with a flexible, hinged joint between the manubrium and dens that bends (up to 90^°^) during their ultrafast leap (Fig. 1D, E), whereas others have a similarly long but much stiffer manubrium-dens joint that bends only slightly (*12*). Certain groups possess comparatively short furcas and still achieve effective escape jumps (*6*). Confocal imaging reveals resilin-rich material at the manubrium–dens joint, especially in species with visibly bending furcas (Fig. 1D,E), suggesting a compliant distal articulation in some but not all springtails. Inspired by this natural diversity, we combine comparative morphometrics, high-speed kinematics and a mathematical model with a centimeter-scale jumping robot (robo-springtail) that can be equipped with furca-like appendages (robo-furca) spanning long-compliant, long-rigid and short morphologies (Fig. 1F–I). We use this joint organism–robot approach to test how furca length and distal compliance influence takeoff angle, body rotation, and jump direction, and to determine whether a single compliant distal hinge can passively stabilize insect-scale jumping robots and maintain performance on uneven and deformable substrates (Fig. 1J–M).

## Results

### Bimodal furca length and distal hinge flexibility define three springtail morphotypes

We quantified body and furca morphology from an online photo dataset of soil- and leaf litter-dwelling springtails collected in the southeastern United States (*20*). The original dataset contained 1,684 images; after filtering for lateral views with the furca fully extended, we retained 552 individuals spanning 15 families across all four springtail orders (Fig. 2A-D; Table S2). For these individuals, we measured the three furca segments (manubrium, dens, and mucro), normalized total furca length by body length (BL), and analyzed the distribution of relative furca length. A Gaussian mixture model (GMM) revealed two modes separated by the intersection of the fitted Gaussian components at 0.25 BL (log-likelihood = 332.5, df = 5, BIC = 633.3; Table S3, S19). The short-furca mode (28% of samples, n=159) was centered at 0.17 BL and long-furca mode (72%, n=393) at 0.48 BL (Fig. 2C). Short-furca individuals were primarily in the order Poduromorpha, whereas long-furca individuals were primarily from the globular-bodied orders Symphypleona and Neelipleona; Entomobryomorpha spanned both morphotypes (Fig. 2A, D). Within each mode, relative furca length scaled with negative allometry (Peak 1 slope = 0.18, *p* < 0.001; Peak 2 slope=0.42, *p* < 0.001; Table S4). This bimodality is consistent with the commonly recognized division between short- and long-furca springtails reported in the natural history literature, which is often associated with more edaphic versus more surface/litter habitats (*6, 13*).

**Figure 2:**
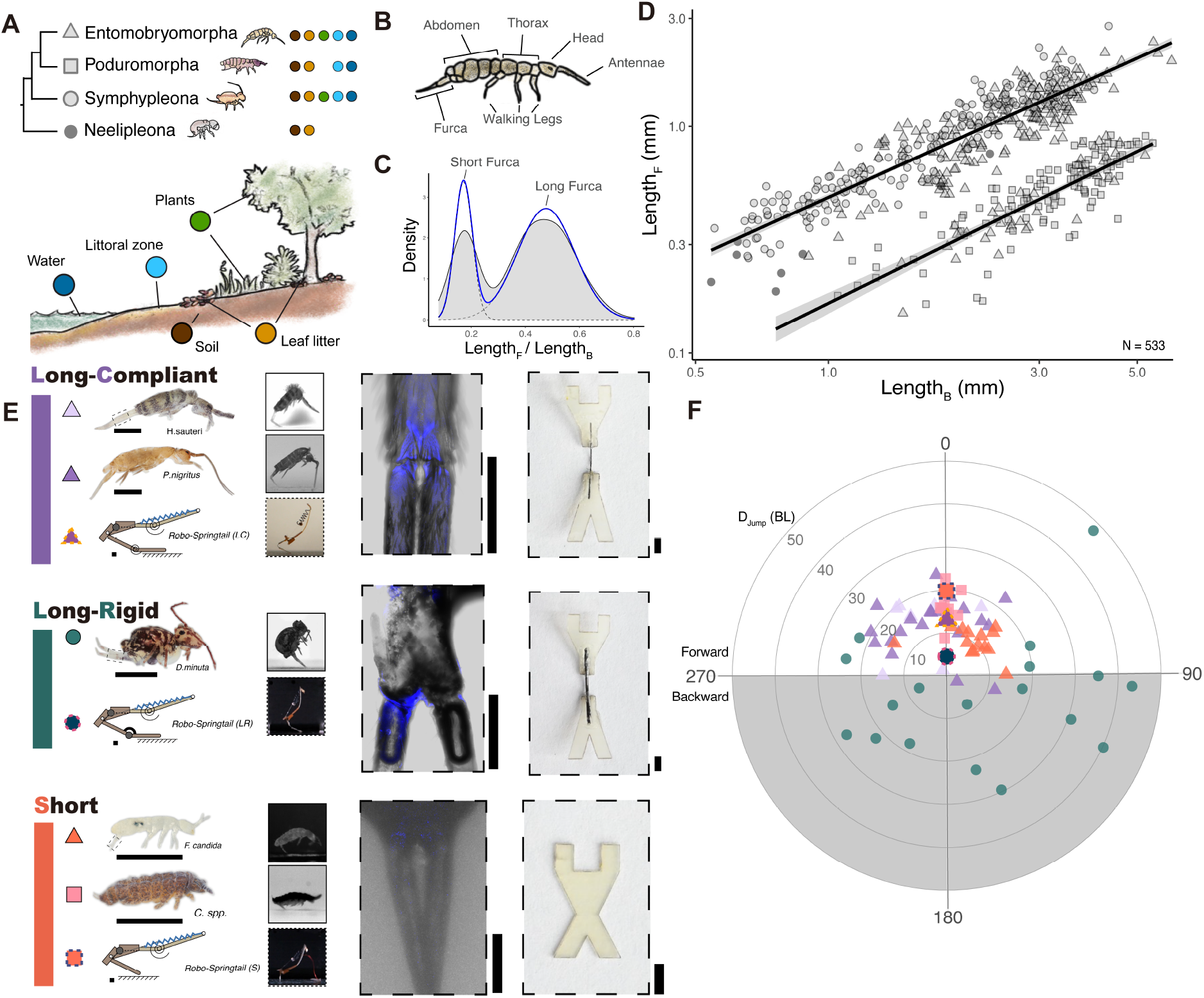
Springtail furca diversity reveals three morphotypes with bi-directional jumps. (A) Phylogenetic tree of the four springtail orders. Colored icons indicate representative environments reported for taxa within each order. (B) Springtail body plan with labeled structures. (C) Relative furca length across n = 552 individuals shows a bimodal distribution; the blue curve shows a Gaussian mixture model fit. Dashed lines mark the relative furca lengths used in the robo-springtails. Scaling of furca length with body length across orders. Points represent individual animals, and black lines show separate allometric fits for the two peaks in (C). (E) Five focal species grouped into three morphotypes alongside corresponding synthetic analogs. High-speed snapshots show peak furca posture during takeoff. Insets show confocal images of the manubrium-dens joint; the fluorescent blue indicates resilin-rich regions. The corresponding robo-furca uses a nitinol wire as an elastic hinge. (F) Escape-jump trajectories (polar plot; radius = jump distance in BL, angle = direction) for the five species and robo-springtail. Point color corresponds with species groups in panel E. The long-rigid morphotype shows predominantly backward-directed jumps, whereas the long-compliant and short morphotypes jump forward.

High-speed videos revealed that, among springtails with relatively long furcas, species differ markedly in the flexibility of the manubrium-dens joint: in some, the joint bends substantially during takeoff (Fig. 2E), whereas in others it remains completely rigid, consistent with previous descriptions of stiff furcas in some taxa (*12*). To examine whether joint material composition differs across these morphotypes, we used confocal light scanning microscopy (CLSM) to image the manubrium–dens articulation in representative species and detected resilin at the joint. Resilin is an elastomeric protein in arthropod cuticle that is often enriched in structures undergoing repeated deformation and rapid elastic recoil (*21, 22*). Resilin signal was present in all imaged specimens, with more prominent resilin-rich regions in species whose manubrium-dens joint visibly bends during the jump (Fig. 2E). This localization is consistent with an elastic hinge that experiences repeated strain during jumping and motivates treating the manubrium-dens joint as a compliant element in our subsequent kinematic and modeling analyses.

To link furca design to jumping trajectories, we selected five species representing three springtail orders and the three furca morphotypes (long-compliant, long-rigid, and short; Table S5). The long-compliant morphotype (lc; *P. nigritus* and *H. sauteri*) has a long furca that bends markedly at the manubrium-dens joint during takeoff, whereas the long-rigid morphotype (lr; *D. minuta*) has a similarly long furca with little visible joint bending (Fig. 2E). The short morphotype (s; *F. candida* and *C. spp*.) has an extremely small furca, and internal joint bending could not be resolved in our lateral views.

From top-view recordings we measured jump distances and directions (n = 72 trials across five species; Table S5; Fig. 2F). Jump distances overlapped broadly across species (ANOVA *p* = 0.077; Table S6), with only modest pairwise differences. However, jump direction depended strongly on morphotype: the long-rigid species *D. minuta* predominantly jumped backward (≈ 150^°^, n = 20), whereas both long-compliant species and both short-furca species jumped forward (Tukey HSD, Watson–Williams tests; Table S5–S7). Thus, either shortening the furca or introducing a bending distal joint is associated with forward-directed escape jumps, a pattern we mirror in the three robo-furca designs shown in Fig. 2E.

### Jointed long furcas lengthen takeoff and reduce body rotation in springtail jumps

In springtail jumps, the furca first releases and the manubrium and dens rotate rapidly away from the body until the mucro, the distal tip of the furca, contacts the substrate (Fig. 3A-C). From high-speed videos, we measured the takeoff duration, defined as the time from furca release to the body leaving substrate, for each species (Table S8). Long-compliant springtails showed the longest takeoff durations (3.7 ± 1.1 ms), whereas long-rigid and short-furca springtails took off much more quickly (1.7 ± 0.1 and 1.1 ± 0.03 ms, respectively; Fig. 3D). As the furca pushes against the ground, the body pitches upward: the anterior region lifts first and the rest of the body follows (Fig. 3A-C and Fig. S1; Movie S1). Long-rigid springtails reached much higher peak angular velocities (100.8 ± 22.3 deg/ms) than either short-furca species (21.9 ± 18.9 deg/ms) or long-compliant species (22.9 ± 8.9 deg/ms) (Fig. 3F; Table S8).

**Figure 3:**
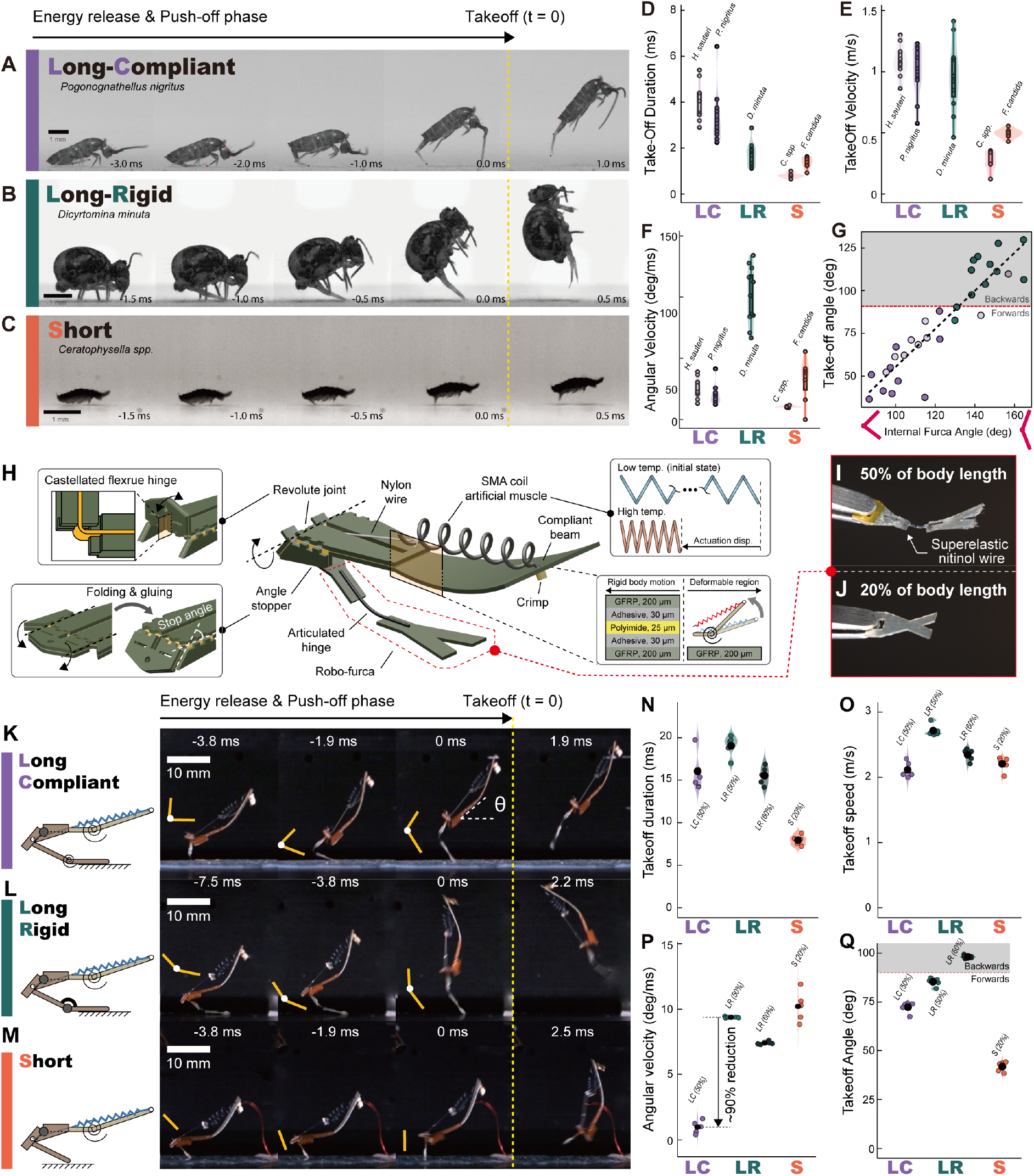
Furca compliance trades takeoff speed for reduced rotation and more forward takeoff in springtails and robots. (A-C) Representative ultrafast jumps of long-compliant (*P. nigritus*), long-rigid (*D. minuta*), and short (*C. spp*.) springtails. (D-F) Take-off duration, takeoff speed, and body angular velocity at takeoff across species and morphotypes (points: individual means; black circles: species means; sample sizes in Table S8). (G) Takeoff angle versus internal furca bend angle, where the red dashed line marks 90^°^, gray shading indicates backward jumps. (H) Springtail-inspired jumping robot architecture and its components. (I) Long robo-furca (50% of BL) (J) Short robo-furca (20% of BL). (K-M) Robo-springtail schematics and push-off snapshots for the three robo-furca configurations. (N-Q) Robot takeoff duration, takeoff speed, average angular velocity, and takeoff angle (n=5 trials per robo-morphotype; black circles: means).

Takeoff angle varies systematically across morphotypes (one-way ANOVA, *p* < 0.001; Fig. 3G; Table S9). Long-rigid springtails took off at higher angles (109.5 ± 14.3^°^), producing backward jumps, whereas springtails in the long-compliant and short-furca morphotypes took off at lower angles that produced forward jumps (Fig. 3G). Within the long-furca species, we measure the internal manubrium–dens angle at maximum bending and find that long-compliant springtails bent the joint more (smaller internal angles) than the long-rigid species, whose joint remained closer to straight. We fit a linear mixed-effects model relating takeoff angle to maximum internal furca angle for the long-furca species and found a significant positive relationship (estimate = 0.88, s.e. = 0.125, *t* (23.67) = 7.09, *p* = 2.7 × 10^−7^; Tables S11). This analysis supports the pattern that more strongly bent furcas (smaller internal angles) are associated with lower, forward-directed takeoff angles, whereas straighter furcas (larger internal angles) are associated with higher, more backward trajectories.

### Mathematical model quantifies the effects of furca length and joint stiffness on springtail jumps

To test whether furca length and distal stiffness are sufficient to explain the observed kinematics, we built a planar Newtonian dynamics model of a segmented springtail with a body and a two-link furca (see SI for details; Fig. S3). For each morphotype, we prescribed the measured body and furca lengths and varied the ratio of torsional stiffness between the distal furca joint and the base joint. With morphotype-specific parameter sets, the model reproduced organismal takeoff duration, translational velocity, angular velocity, and takeoff angle; across all 4 metrics, model predictions lie within 1–10% of the corresponding organismal means (Figs. S2-S3).

The model also clarifies why morphotypes differ in jumping performance (Figs. S3I–L). In the simulations, short furcas produce brief contact times and lower takeoff speeds, whereas long furcas produce longer jump durations. Geometric and dynamic scaling predict that larger jumpers rotate more than smaller ones even when the takeoff angle and speed are similar. Finally, when we sweep joint stiffness ratio in the long-furca regime, angular velocity varies non-monotonically with stiffness: very flexible and very stiff joints both suppress rotation relative to intermediate stiffness values (Fig. S3K). Long-compliant species such as *H. sauteri* and *P. nigritus* occupy the compliant, low-rotation region, whereas the long-rigid morphotype lies toward the stiff end of this landscape, consistent with their contrasting angular velocities in vivo. This length–stiffness map, therefore, provides a mechanistic link between organismal kinematics and the furca designs we implement in our jumping robots next.

### Compliant furca hinge stabilizes robo-springtail jumps on flat ground

The results from our mathematical models and organismal experiments indicate that jumping performance (speed and takeoff angle) and aerial stability (body rotation) depend on the length and compliance of the springtail furca. To assess whether the same principles hold for insect-scale robots, we built a 20-mm, 84-mg robo-springtail with an interchangeable furca-like jumping appendage (robo-furca) as shown in Fig. 3H-J. The robo-furca is a laser-cut composite structure with a castellated flexure hinge and, for the compliant configuration, a superelastic nitinol wire forming the distal joint (Fig. 3H-J; fabrication details in Methods and Fig. S4). To evaluate the three morphotypes, we used robo-furca configurations that matched a short furca (20% of BL) and a long furca (50% and 60% of BL) with either a rigid or an elastic hinge (Figure 3K-M; Movie S2; Fig. S5-7).

We tracked the robot’s body posture and kinematics during jumps and found that with either the short furca or the long-rigid furca, the robot rotates rapidly because the line of action of the ground reaction force lies far from the body’s center of mass (Fig. 3L,M). When we equip the longer furca with a compliant hinge, the robot instead settles to a stable body angle of ∼ 40^°^ as the hinge deforms during push-off, redirecting the ground reaction force closer to the center of mass (Fig. 3K). Robots with longer furcas have longer takeoff durations than the short-furca ones (Fig. 3N). Among the long-furca configurations, the rigid hinge yields a longer takeoff duration than the compliant hinge version, consistent with our mathematical model (Fig. 3L).

Takeoff velocities also parallel the biological data (Fig. 3D-G). The long-rigid robot reaches the highest takeoff speed (2.74 ± 0.09 m/s and 2.37 ± 0.09 m/s at 50% and 60% of BL, respectively), followed by the short-furca robot (2.23 ± 0.12 m/s) and the long-compliant robot (2.13 ± 0.12 m/s) (Fig. 3O). Takeoff angles increase from the short-furca robot (41.75 ± 2.56°) to the long-compliant robot (72.42 ± 2.84°) and are largest for the long-rigid robot (85.35 ± 2.08°and 98.09 ± 0.82°at 50% and 60% of BL, respectively), mirroring patterns in springtails.

We next measured the full jumping trajectory and the robot’s orientation in mid-air to compute angular velocity (Fig. 3N-Q; Fig. S7; Movie S2). The compliant-hinge configuration clearly out-performs the rigid configurations: the angular velocities of the short-furca and long-rigid (50% and 60% of BL) configurations are 10.21 ± 1.18 deg/ms, 9.38 ± 0.05 deg/ms, and 7.38 ± 0.12 deg/ms, respectively, whereas the robot with the compliant hinge has an angular velocity of 0.98 ± 0.49 deg/ms, ∼ 90% lower than that of the long-rigid configuration (Fig. 3P).

The robo-springtail experiments therefore show that different robo-furca morphotypes yield distinct jumping behaviors. A short furca enables rapid push-off, allowing the robot to reach high takeoff speeds with the shortest takeoff duration. A long, rigid furca produces the highest takeoff velocity in a nearly vertical direction but generates large body rotations. By contrast, a long furca with a compliant hinge provides passive posture control by substantially reducing body rotation: its takeoff velocity is only 4.2% lower than that of the short-furca robot and 22.1% lower than that of the long-rigid robot, yet this trade-off yields aerially stable, forward-directed jumps on flat ground (Fig. S7B).

### Compliant hinge maintains takeoff speed and limits rotation on rough terrain

Biological jumpers such as springtails operate on complex natural substrates (soil, plants, leaf litter, water) that are uneven and deformable, unlike the flat, rigid surfaces typical of jumping-robot experiments (*23*). To explore whether a passively compliant furca joint improves performance on more realistic terrain, we performed additional experiments on uneven and gravel substrates using both the long-rigid and long-compliant robo-springtails.

To quantify substrate roughness, we fabricated surfaces with periodic pairs of semicircles of radius *R*, arranged symmetrically above and below a baseline (Fig. 4A,B; Fig. S8). We oriented the robot so that its jump direction aligned with surface waveform. During push-off, the rigid-hinge robot pushes off through a single contact point, whereas, the compliant hinge deforms strongly on rough surfaces, effectively “spreading” into the valleys between raised sections (Fig. 4C,D; Movie S3). This deformation helps the furca maintain ground-reaction forces even when local slippage occurs.

**Figure 4:**
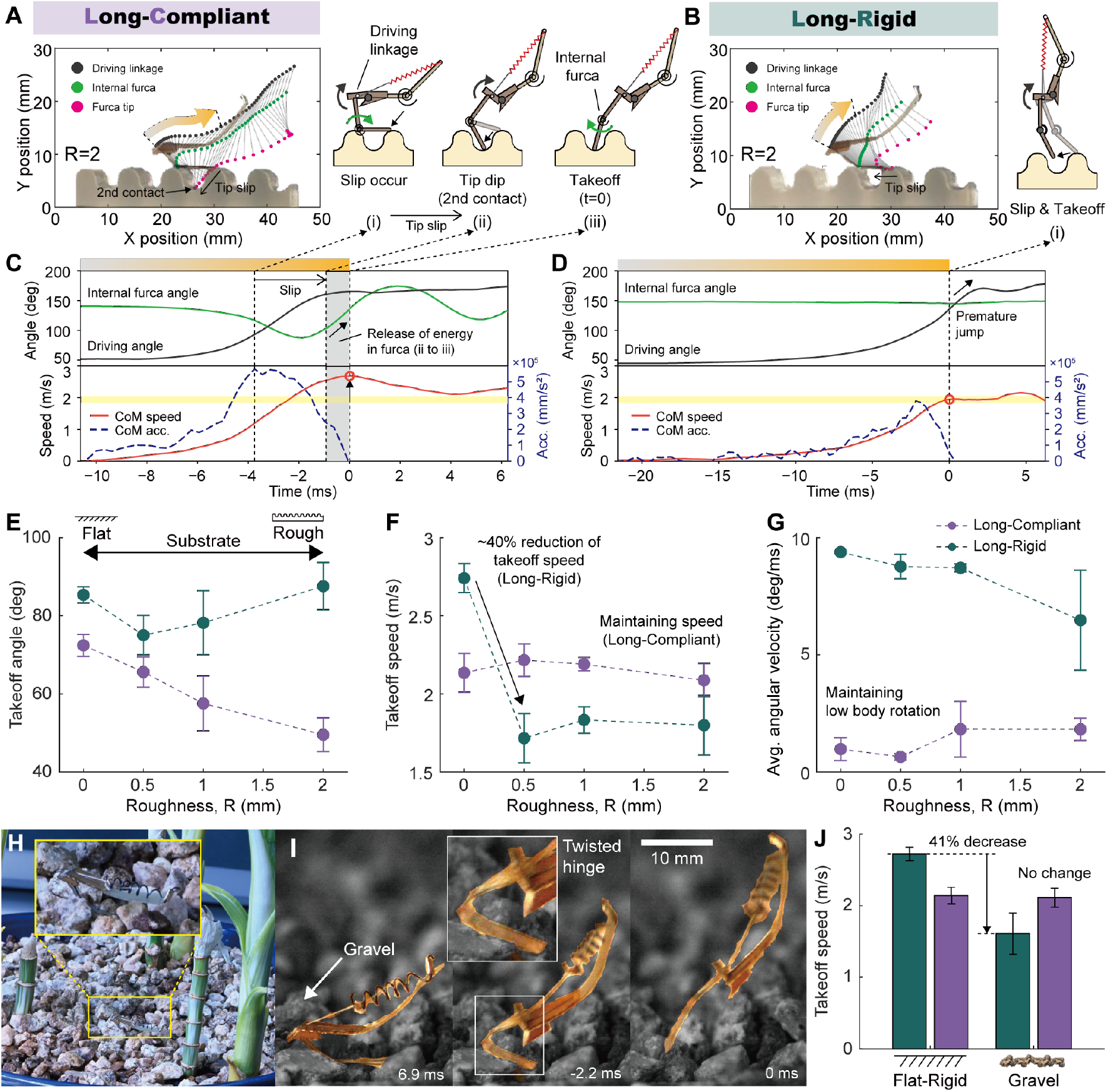
A compliant robo-furca hinge maintains takeoff speed and limits body rotation on rough terrain. (A-B) Furca-tip trajectories and key contact events during push-off for the long-compliant and long-rigid configurations on a rough surface with roughness amplitude *R* = 2 mm. (C, D) Time series of driving-linkage angle, internal robo-furca angle, center-of-mass speed, and acceleration during push-off for the long-compliant and long-rigid configurations. (E-G) Takeoff angle, takeoff speed, and mean body angular velocity as functions of surface roughness *R* (mean ± s.d., *N* = 5 trials per condition). (H,I) Naturalistic setting and representative jump sequence on gravel highlighting hinge deformation (inset). (J) Comparison of takeoff speed on a flat rigid substrate and on gravel (mean ± s.d.; flat *N* = 5, gravel *N* = 3).

With a rigid hinge, takeoff angle increases with roughness, as if the robot were jumping uphill while maintaining the initial contact point between furca and substrate (Fig. 4E). In contrast, the compliant-hinge robot shows decreasing takeoff angle with increasing roughness, consistent with the hinge deforming “downhill” into the rough terrain (Fig. 4E). On rough surfaces, the rigid-hinge robot’s takeoff velocity drops by more than 33% relative to flat ground, whereas the compliant-hinge robot shows little loss in its takeoff speed (Fig. 4F; Fig. S9-S10; Movie S3). The long-compliant robo-springtail also shows much lower body rotation, remaining below 1.8 deg/ms across rough surfaces, while the rigid-hinge robot exceeds 6.5 deg/ms (Fig. 4G).

We extended this analysis to natural terrain by testing on gravel substrates with a range of stone sizes, which introduce more variable and unpredictable contacts than the printed surfaces (Fig. 4H; Movie S4). Gaps between stones reduce contact-force reliability because the gravel can shift under load (Fig. 4I and Fig. S8C). Even so, the compliant-hinge robot adapts and largely preserves its jumping performance, whereas, the rigid-hinge robot shows a 41% decrease in takeoff speed compared to flat ground (Fig. 4J).

### Compliant hinge mitigates performance loss on deformable substrates

Beyond roughness, natural substrates can deform under push-off forces and dissipate energy, reducing takeoff speed and stability (*14, 15, 23–27*). To examine whether a compliant robo-furca joint mitigates these losses, we fabricated springboards with tunable bending stiffness by varying beam lengths *L*_*b*_ (Figs. 5A,B; Fig. S8B; Movie S5).

**Figure 5:**
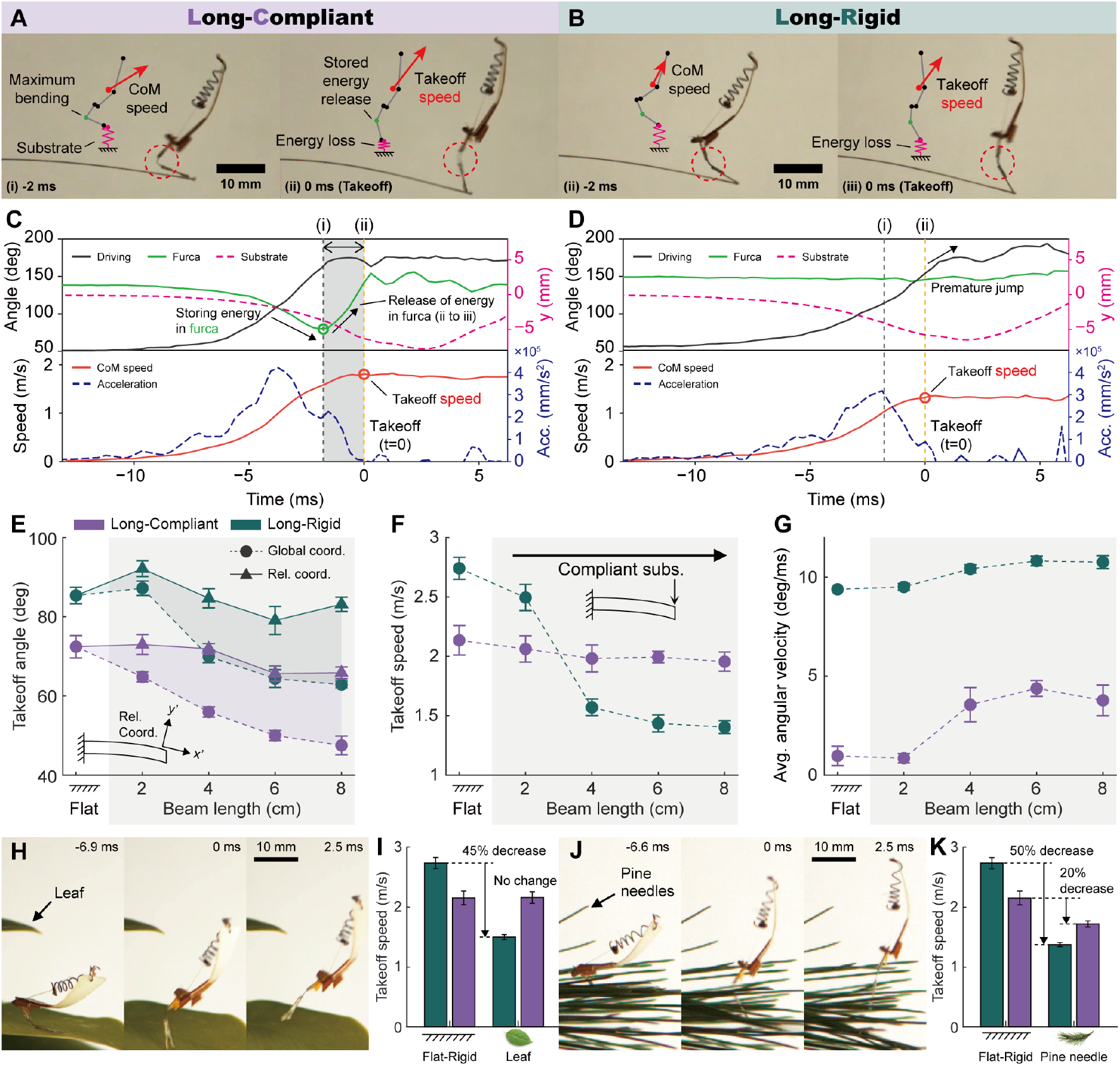
Compliant hinge preserves robo-springtail takeoff performance on flexible beams, leaves, and pine needles. (A,B) Representative high-speed snapshots of takeoff for robots with long-compliant and long-rigid robo-furcas. (C,D) Time series of driving-linkage angle, internal robo-furca angle, substrate deflection, center-of-mass speed, and acceleration during takeoff for both configurations. (E-G) Takeoff angle (reported in the global frame and a substrate-attached frame), takeoff speed, and mean body angular velocity as a function of substrate compliance (beam length). Points show mean ± s.d. (N=5). (H,J) High-speed sequences of the compliant-hinge robot taking off from leaves and pine needles. (I,K) Takeoff speed on leaf and pine needles compared to flat ground for both configurations (flat: N=5; natural substrate: N=3; mean ± s.d.).

With a compliant hinge, the robot shows near-constant takeoff speed across substrate compliances despite substantial beam deflection. Center-of-mass acceleration shows a secondary peak that coincides with hinge recoil (recovery of the internal robo-furca angle from its maximum bend), and this impulse becomes more pronounced on more compliant beams (Fig. 5C; Fig. S11-S12). In contrast, the rigid-hinge robot shows no comparable hinge-recoil impulse, and its takeoff speed decreases as substrate compliance increases, consistent with energy loss to substrate deformation (Fig. 5D,F). On longer beams, the rigid-hinge robot often takes off before the substrate rebounds, reducing the impulse available from substrate recoil.

In the global frame, takeoff angle decreases with increasing beam length for both hinge types because the robot tilts with the deflecting substrate. In a substrate-attached (relative) frame, takeoff angle varies much less with beam length and remains lower for the compliant hinge than for the rigid hinge (mean 69.1^°^ vs 84.7^°^, averaged across compliant beams; Fig. 5E).

Substrate compliance substantially degrades the rigid-hinge robot: takeoff speed drops by as much as 43% on a 4 cm beam relative to flat ground, whereas the compliant-hinge robot sustains an approximately constant takeoff speed of ∼ 2 m/s over the same range (Figs. 5F,G). Angular velocity increases with substrate compliance for both hinge types, but the compliant-hinge robot exhibits 70% fewer body rotations than the rigid-hinge robot (Fig. 5G).

Lastly, to test more natural compliant substrates, we repeated these experiments on leaves and pine needles (Movie S6). On leaves, the compliant-hinge robot shows little loss in takeoff performance, whereas the rigid-hinge robot shows an 45% decrease in takeoff speed relative to flat ground (Fig. 5H,I). On pine needles, which deflect more readily, the rigid-hinge robot loses 50% of its takeoff speed, while the compliant-hinge robot remains relatively stable with a smaller 20% speed reduction (Fig. 5J,K).

## Discussion

### Summary

Across springtails, furca length and joint compliance map onto distinct jump strategies. Long-rigid springtails take off rapidly but rotate strongly and preferentially launch at high angles (backward-jumps), whereas long-compliant springtails bend a resilin-rich manubrium–dens hinge to produce longer takeoff durations, forward-directed trajectories, and reduced rotation. A mathematical model shows that varying furca length and joint stiffness can account for these kinematic trade-offs. Implementing a compliant robo-furca in a 20-mm, 84-mg robo-springtail reduces body rotation by ∼90% and preserves takeoff performance across irregular and yielding substrates, supporting hinge compliance as a passive route to directional control and postural stability at the insect scale.

### Ecological trade-offs in furca design

Springtails are among the most abundant soil animals and occur globally from the tropics to polar regions. Global syntheses estimate ∼ 2×10^18^ individuals in soils worldwide, with a total biomass of ∼ 27.5 Mt C, roughly threefold higher than the carbon biomass of all wild terrestrial vertebrates (*7*). Springtails also sit on a steep ecological gradient, from euedaphic taxa moving through soil pore networks to epedaphic and plant-associated taxa navigating exposed, heterogeneous surfaces. Classical life-form frameworks link this vertical stratification to suites of morphological traits, including furca development (*28*). Our bimodal distribution of relative furca length aligns with this view: reduced-furca phenotypes are common in soil-associated taxa, whereas long-furca phenotypes are common among surface-active taxa where dispersal and rapid relocation can matter in patchy or risky habitats (*6, 29*).

Within the long-furca regime, our results add a second functional axis, distal hinge compliance, that shifts performance away from maximizing initial impulse and toward controlling takeoff direction and body rotation during an ultrafast launch. This trade-off may matter on substrates such as leaf litter, snow, or at air-water interfaces, where a high-angle, high-rotation takeoff could increase landing variability and escape from predators, while a slightly slower but more predictable forward launch could support repeated jumps and navigation. Semiaquatic springtails illustrate how takeoff direction and landing success can couple tightly to contact mechanics and body posture at an interface (*11*). The resilin-rich material we observe at the manubrium-dens region provides one plausible structural route for embedding this control in the furca itself. What remains unclear is whether hinge compliance yields a measurable fitness advantage in the field, and how often this compliance appears across springtail lineages or changes across ontogeny.

### Mechanical intelligence at the contact interface in ultrafast jumpers

Springtail jumps use latch-mediated spring actuation (LaMSA) (*12,30*). LaMSA solves the ultrafast power-delivery problem, but it compresses control into the milliseconds after latch release. During that brief push-off, there is little opportunity for feedback to correct posture or compensate for unexpected contact mid-event. Latches and linkages, therefore, do more than trigger release; they shape the timing and direction of the impulse delivered to the environment, and in turn the trajectory and stability of the movement (*31*). Here, we find that furca compliance can shape the takeoff impulse at ground contact, where friction, roughness, and substrate deformation vary the most.

Furca compliance can thus be viewed as a ‘mechanical filter’ that reduces sensitivity to substrate-dependent contact. Furca compliance changes contact geometry during push-off and can effectively shift the ground-reaction line of action relative to the center of mass, reducing body rotation without requiring online sensing or computation during the launch. This places compliance where it matters most for repeated performance: not just in the energy-storage element that powers the jump, but also at the contact interface that transmits that energy into the physical world. Similar intrinsic mechanical stabilization appears in legged locomotion on irregular terrain (*32, 33*) and in robotics, where series elasticity improves tolerance to contact uncertainty in rapid physical interactions (*34*). A key open question is whether springtails inherit hinge stiffness passively from cuticle architecture (e.g., resilin-enriched regions) or whether they can tune the joint before release through posture, muscle loading, or pre-release furca motions (*35*).

### Control challenges for insect-scale jumping robots

Passive mechanisms are attractive in insect-scale robots because they add function without adding active actuators, sensing, or power draw. In mass- and power-limited platforms, passive elements already improve performance, from energy-efficient locomotion (*18*) to wing deployment and retraction (*36*). Jumping in real terrain is harder because the robot has to (i) achieve terrain-adaptive takeoff (ii) set takeoff direction, (iii) stabilize in flight, (iv) land in a recoverable pose, and (v) reload for the next jump (Fig. 1C).

Our findings suggest that furca compliance improves takeoff, the first step in the jump cycle. The compliant robo-furca reduces angular impulse and stabilizes launches across naturalistic terrain without additional sensing or active control. This complements, rather than replaces, strategies that target other phases, including designs that shape takeoff through posture or morphology (*37*) and passive mechanisms that bias aerial self-righting and controlled landing (*11, 38*). A remaining engineering challenge is to co-design takeoff adaptability, landing biasing, and rapid reloading, so that an insect-scale robot can close the full cycle and jump repeatedly and autonomously in natural, obstacle-filled terrain (Fig. 1C).

### Terrain adaptability on compliant substrates

Substrate compliance complicates jumping because the ground becomes part of the launch dynamics. Work done during push-off can be diverted into substrate strain energy and only returned on the substrate recoil timescale. Across biological and engineered jumpers spanning orders of magnitude in mass, takeoff speed typically decreases as substrate compliance increases (Fig. 6B; Table S12-13). Takeoff duration also varies strongly with scale (from ∼1 ms to > 100 ms; Fig. 6A), which sets how much time a system has to adjust posture, limb mechanics, or the timing during push-off.

**Figure 6:**
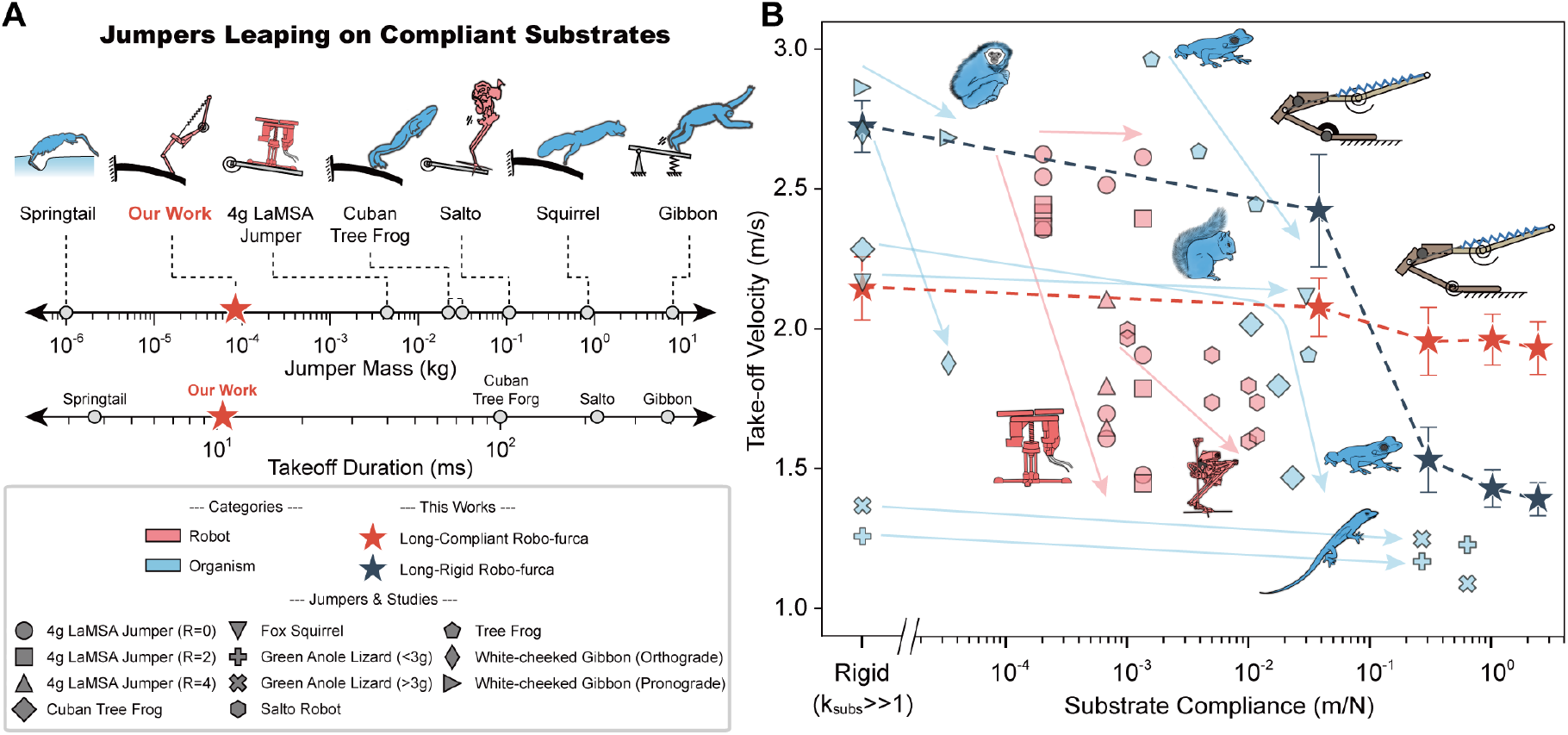
At the low-mass end of terrain-adaptable jumpers, a compliant hinge maintains millisecond takeoff on deformable substrates. (A) Representative biological and robotic jumpers tested on compliant substrates, positioned by body mass and takeoff duration (Table S13). Stars denote the robo-springtail configurations used in this work. (B) Takeoff speed versus substrate compliance for biological and robotic jumpers. Across systems, takeoff speed typically decreases as substrate compliance increases. Our compliant-hinge robo-furca shows a flatter speed-compliance trend than the rigid-hinge configuration. Data sources in Table S13.

Larger vertebrates and motor-driven robots can partially mitigate compliance-induced losses by adjusting posture and effective limb stiffness and, in some cases, timing takeoff relative to substrate recoil (*26, 32, 39, 40*). Even with active adjustment, small changes in substrate stiffness or effective mass can shift recoil phase and reduce the impulse available at takeoff (*25,40,41*). Robotic platforms at these scales can also demonstrate controlled takeoff and landing behaviors through active control strategies (*16, 17*).

At insect scales, takeoff unfolds over milliseconds, which makes it difficult to time takeoff to match substrate recoil. Our data show that distal hinge compliance reduces this sensitivity, since the hinge bends as the substrate deflects and then recoils, adding a secondary push-off impulse. As a result, the compliant-hinge configuration shows a weak dependence of takeoff speed on substrate compliance while also limiting body rotation (Fig. 6B). In contrast, the rigid-hinge configuration shows a much stronger drop in takeoff speed as compliance increases. Compared with prior robots tested across compliant substrates, our platform operates at more than an order of magnitude lower mass, placing it at the low-mass end of terrain-adaptable jumpers (Table S13).

Future work should test how well this strategy generalizes to substrates that combine compliance with other field-relevant complications (e.g., water surfaces, snow, and granular media), and whether it can expand the performance envelope of small robots. Springtails navigate these environments routinely, suggesting that additional design rules remain to be extracted from insects and other small-scale jumpers. More generally, our work adds to a growing set of examples in which geometry, topology, and material physics shoulders part of the control burden in small-scale robots, including instability-driven jumping and other forms of physical intelligence (*42–44*).

## Acknowledgments

We thank the Bhamla Lab members for useful discussion and feedback. We thank Anthony Senft for his help in data collection. We thank Drs. Felipe Soto-Adames and Rosanna Giordano for expert guidance on springtail specimen identification. We also thank Dr. Michael Caterino for allowing us to use his image database of soil-dwelling arthropods.

## Author contributions

J.H., B.K., J.-S.K., and S.B. conceptualized the research. J.H., A.S., and T.T. designed the animals’ experimental framework. B.K. and J.-S.K. contributed to the design of the jumping appendages and jumping robot. B.K. and J.-S.K. developed the robot’s experimental design. H.K. conceived the mathematical model of the jumpers. S.B. and J.-S.K. provided funding for the project and supervised the research. J.H., B.K., H.K., A.S., T.T., J.-S.K., and S.B. contributed to writing, reviewing, and editing the manuscript.

## Data and materials availability

All data are available in the manuscript or the supplementary materials.

## Supplementary materials

Materials and Methods

Supplementary Text

Figs. S1 to S13

Tables S1 to S15

References *(7-66)*

Movies S1 to S6

