## Supplementary Materials for "Springtail-inspired compliant hinge enables terrain-adaptable takeoff in insect-scale robots"

**This PDF file includes:**

Materials and Methods
Supplementary Text
Figures S1 to S13
Tables S1 to S15
Captions for Movies S1 to S6

**Other Supplementary Materials for this manuscript:**

Movies S1 to S6

### Materials and Methods

#### Comparative Morphology

We used descriptions of springtail families in *Biology of the Springtails* (*6*) to look at the presence of a furca and the preferred habitat of the family. Springtail families vary in the number of species and habitat preference, so this is a rough approximation of family preference across springtails. We then placed the information on a phylogenetic tree of springtails pruned to include only the order level for which we had information about their furca and habitat preferences (Fig. 2A).

We examined morphological variation in springtail furcas using an online photo-database of soil-dwelling organisms collected by Dr. Michael Caterino (*20*). Specimens were collected by sifting the leaf litter and top layers of soil, placing the soil samples on Berlese funnels, and collecting the organisms that filtered through the funnels in 70% EtOH. Images were taken from Berlese funnel extractions of leaf litter across the southeastern United States and photographed with a Canon EOS 6D and MP-E65mm lens. Of 1,684 images, we retained 552 lateral-view images of individuals with the furca fully extended. Images of specimens were identified to family with the assistance of experts (Felipe Soto-Adames, Curator of Arthropods, Florida Department of Agriculture, and Rosanna Giordano, Florida International University).

After filtering, we measured body length, manubrium, dens, and mucro using calibrated scale bars (0.1 mm). Furca length was calculated as the sum of the three sections and normalized to body length. To assess distribution patterns, we fit a Gaussian Mixture Model (GMM) using the `mclust` package in RStudio (v2024.09.1+394) and tested for multimodality with Hartigan's dip test (Table S14). Individuals were grouped according to the intersection point (0.25 body lengths) between Gaussian components. Scaling relationships were analyzed with linear regressions within each group. Results for statistical tests can be found in the results section and the supplemental information.

#### Study Species

We selected five species representing three springtail orders and three morphotypes: long-compliant (*Pogonognathellus nigritus*, *Homidia sauteri*), short (*Folsomia candida*, *Ceratophysella* spp.), and long-rigid (*Dicyrtomina minuta*). *P. nigritus* and *D. minuta* were collected by live-sifting leaf litter through a metal mesh in the author's (AAS) residential neighborhood in Raleigh, North Carolina. *H. sauteri* was collected from oak litter on the Georgia Institute of Technology campus in a similar manner. *F. candida* and *Ceratophysella* spp. were purchased from Springtails.us. Species identifications were confirmed using taxonomic keys (*45*).

#### Directional Jumps

We elicited escape jumps by lightly touching the abdomen with a brush or pin and filmed trajectories from above. *P. nigritus* (N=20 jumps, 20 individuals) and *D. minuta* (N=20 jumps, 20 individuals, from a previously published dataset (*12*)) were filmed with a Phantom Miro LC321S camera (Laowa 60mm lens, 1280×800 px, 2,948 fps). *H. sauteri* (N=22 jumps, 10 individuals), *F. candida* (N=24 jumps, 16 individuals), and *C. spp*. (N=29 jumps, 10 individuals) were filmed at 1,000 fps with a Photron FASTCAM Mini and Nikon 70–200mm f/2.8 lens (1024×1024 px). Calibration was performed with a 0.5 mm scale in view. Jumps obstructed by the probe were excluded.

Jump distance and direction were extracted from takeoff to first substrate contact, measured relative to initial body orientation. Distance were compared across species with ANOVA followed by Tukey's HSD. Jump angles were analyzed with a Rayleigh test for uniformity and Watson–Williams tests for circular means, including pairwise comparisons, using the `circular` package in R. All data visualization was performed with `ggplot2`.

#### Furca Morphology

We examined furca morphology (manubrium, dens, and mucro) using confocal laser scanning microscopy (CLSM) for *D. minuta*, *H. sauteri*, and *F. candida*, focusing on resilin at the manubrium–dens joint. Unless noted, individuals were preserved in 70% EtOH prior to preparation (Fig. 2E). To visualize resilin, we dissected the furca by cutting at the manubrium base under 70% EtOH. Slides were prepared with transparent self-adhesive reinforcement rings to create a shallow chamber matched to specimen thickness. Chambers were filled with glycerine, the furca was positioned, and a coverslip was sealed with cyanoacrylate adhesive.

Samles were imaged with a Zeiss LSM710 confocal microscope. Z-stack images were acquired through the depth of the sample and subsequently stitched together using ImageJ (*46*). Maximum intensity projections (or 3D reconstructions when applicable) were generated for analysis. Consistent with established practice for resilin autofluorescence, we excited at 405 nm and detected emission at 420–480 nm; this band reliably visualizes resilin with modern CLSM systems that lack UV lasers and supports material-contrast mapping when combined with brightfield overlays (*21*). Laser power, detector gain, and offset were adjusted to avoid saturation and minimize bleaching (*21*). High-resolution images of the body and furca were captured for all species used in the study, using a Canon EOS 6D and an MP-E 65 mm lens.

#### High-Speed Video Analysis

We filmed lateral takeoffs at ultra–high frame rates to resolve sub-millisecond motion. *Pogonognathellus nigritus* was recorded at $40,322$ fps using a Phantom VEO 1310 with a Canon MP-E 65 mm macro lens (Resolution: $640 \times 480$ px). *Homidia sauteri*, *Folsomia candida*, and *Ceratophysella* spp. were recorded at $40,000$ fps using a Photron SA-Z with the same lens. Scenes were front-lit with a high-intensity LED array and backlit with a diffused LED panel (Neewer CN-160) fitted with a #116 Tough White diffusion filter (Rosco). Takeoff sequences for *Dicyrtomina minuta* were sourced from a previously collected dataset and reanalyzed with the pipeline below. Animals jumped from either a flat acrylic plate or a 3D-printed stage (Bambu Lab X1C, USA) covered with masking tape to provide repeatable friction. A 0.5 mm ruler was used for calibration.

Videos were tracked frame-by-frame in MATLAB (R2018a) using `DLTdv8` (*47*). We digitized six landmarks per frame (Fig. 3A). These were the base of antennae, the head–thorax junction, the abdominal tip, the manubrium base, the manubrium–dens joint, and the mucro tip. Tracking began several frames before release and extended past takeoff. Trials in which the probe or brush visibly impeded motion or obscured landmarks were excluded.

Body planar angle ($\theta_{\text{body}}$) was computed from the vector joining the head–thorax junction and abdominal tip and reported as change from initial orientation. Displacement of the body centroid was measured using the per-frame mean of the three body landmarks (antennae base, head–thorax junction, abdominal tip). External furca rotation was computed with a four-point construction (head–thorax junction, abdominal tip, manubrium base, manubrium–dens joint). We set $0^\circ$ at the pre-release frame and reported subsequent angular displacement between body and furca lines. Internal furca angle (manubrium–dens bend) used a three-point measurement (manubrium base, manubrium–dens joint, dens tip). Because the furca was too small to resolve reliably, the internal angle was not quantified for *F. candida* and *Ceratophysella* spp. jumps.

Displacement and angle trajectories were smoothed by fitting 5th-order polynomials to each time-series data. Velocities and accelerations were obtained from the first and second derivatives of the fits, respectively. Takeoff duration was defined as the interval from furca release to substrate detachment. Takeoff speed was evaluated at detachment. Peak body angular velocity and maximum internal furca angle were extracted from the fitted trajectories.

We compared takeoff angle, takeoff velocity, takeoff duration, peak body angular velocity, and maximum internal furca angle across species. We assessed normality using Shapiro–Wilk tests and used one-way ANOVA with Tukey's HSD when assumptions were met or used Kruskal–Wallis tests with Bonferroni-corrected Wilcoxon post hoc comparisons. Within long-furca species, the relationship between takeoff angle and maximum furca angle was tested using a linear mixed-effects model fit by REML (`lme4`), with takeoff angle as the response, maximum internal furca angle as a fixed effect, and species as a random intercept. For comparison, we also fit simple linear model without random effects and contrasted models by AIC. Analyses and graphics were performed in RStudio using `tidyverse`, `ggplot2`, and `lme4`.

#### Mathematical model

The mathematical model in this manuscript was performed using a custom-made MATLAB program. The program utilized first-order Euler integration of the Newtonian dynamics of a multi-segmented rigid-body in two dimensions. The simulated springtail was considered to have three main degrees of freedom: its $x$ and $y$ planer locations and its body orientation. In addition, the simulated springtail had two internal angles that described the body deformation (Fig. S3A). The mass and moment of inertia were calculated by approximating the body as a cylinder and neglecting the mass of the furca.

In contrast with our previous model (*11*), furca joint kinematics were not prescribed. Instead, the two furca joints (one at the furca base, and the other within the furaca; Fig. S3A) were modeled as linear torsional springs with equilibrium angles $\theta_{1,0} = \theta_{2,0} = 180°$. Joint torques were proportional to angular displacement from equilibrium and coupled to the ground reaction force at the furca tip. Coulomb friction was used to model interaction with the ground.

At each time step, we intially assumed the contact point at the tip of the furca remained fixed.

We calculated the ground reaction force based on the deformation of the springtail (displacement of the two joint angles). If the tangential component of the ground reaction force exceeded what the Coulomb friction limit, the furca slipped and the contact point was recalculated. The solver then entered an iterative routine that calculated a new location of the contact that satisfied all the constraints. When a solution was not viable, the simulated springtail was set to the airborne state with zero ground reaction force.

**Robot fabrication**

The robot was built around the TRC mechanism capable of rapid torque production (*11, 38*). The TRC mechanism comprised a foldable composite body structure actuated by shape-memory alloy (SMA) coil artificial muscles. The body structure was fabricated using the smart composite microstructure (SCM) process (*48*). The five-layer laminate consisted of a facet layer (GFRP, 200 $\mu$m), adhesive (heat-reactive film, 25 $\mu$m), flexure layer (polyimide, 25 $\mu$m), adhesive, and a second facet layer as shown in Fig. S4A. Each layer was laser-cut (Series A, Oxford Lasers Ltd.), and the stack was laminated using a heat press (QM900A, QMESYS). SMA coils were manufactured by winding SMA wire (Dynalloy Co., USA). The wire and core diameters were 250 $\mu$m and 1.5 mm, respectively. The assembled robot body had a body length of 20 mm and a mass of 84 mg. Detailed fabrication of the robot body and actuator is described in (*38*).

Robo-furca jumping appendages were laser-cut (Series A, Oxford Lasers Ltd.) in three configurations (Fig. 3H-J) corresponding to springtail morphotypes: short (20% of BL), long (50% and 60% of BL) with a rigid distal hinge, and long (50% of BL) with a compliant distal hinge. Give the 20 mm robot body length, we set robo-furca lengths to 4 mm (short) and 10 mm (long) (Fig. S4B,C). The short robo-furca was fabricated from a single 200 $\mu$m GFRP sheet. The long robo-furca comprised of two GFRP linkages connected by a hinge made from superelastic Ni-Ti wire (McMaster-Carr). Ni-Ti wire diameters were 250 $\mu$m for the rigid hinge and 100 $\mu$m for the compliant hinge. Each robo-furca was mounted by inserting a notched socket into the robot body (Fig. S4D).

#### Experimental setup of the robot

Robot jumps on flat, rough, and compliant substrates were filmed using a high-speed camera (Phantom MIRO EX4 and MIRO C320, Vision Research, USA). We recorded two sets of videos. To analyze push-off kinematics, we recorded at 3,200 frames per second (Fig. 3K-M; Fig. 4C, D, and J; Fig. 5C, D, I, and K; Fig. S6, S9, and S11). To quantify jump performance (takeoff speed, jumping height and distance, and average angular velocity), full trajectories were recorded at 1,480 frames per second (Fig. 3N-Q; Fig. 4E-G; Fig. 5E-G; Fig. S7, S12, and S13). In both cases, motion was captured in the sagittal plane. The $x$-axis denotes horizontal displacement (forward direction), and the $y$-axis represents the vertical position (height).

To evaluate the effect of robo-furca compliance on uneven terrain, we prepared two rough substrates. The first was a custom-fabricated undulating surface with a defined curvature radius. It was printed with an Objet350 Connex3 (Stratasys Ltd.) 3D-printed using a 1:1 blend of TangoPlus™ and VeroClear™ (Fig. S8A). The second was a gravel surface representing natural terrain, consisting of randomly distributed gravel particles spanning 1–10 mm in size (Fig. S8C).

For compliant terrain, we also prepared two types of substrates. One had mechanically defined properties (flexible beam), and one represented deformable substrates found in naturalistic environments. To tune the stiffness in the engineered flexible substrate, we fabricated a springboard from 200 $\mu$m GFRP sheets using a laser cutter (PLS6.75, Universal Laser Systems, Inc.) (Fig. S8B). Jumping experiments were conducted with beam lengths of 2, 4, 6, and 8 cm. Mechanical properties are listed in Table S15.

#### Analysis of the robot's jumping procedure and performances

We analyzed robot push-off by estimating joint kinematics from zoomed-in 3,200 fps recordings. Joint positions were tracked in the sagittal plane as illustrated in Fig. S13. For compliant substrates, the displacement of the substrate tip was tracked using the same procedure. Joint angles were computed from the tracked positions. The robot's center of mass (CoM) was computed from its kinematic configuration and mass distribution, with procedures and equations described provided in the Supplementary text and Fig. S13. CoM velocity and acceleration profiles were obtained by numerically differentiating the CoM position.

Full jump trajectories was tracked from the instant the vertical ground reaction force reached zero, which corresponds to the detachment of the robo-furca tip from the substrate. Trajectories recorded at 1,480 fps were used to compute takeoff angle, takeoff speed, and average angular velocity (mean rotational velocity over the airborne phase).

### Supplementary Text

#### Calculation of robot's center of mass (CoM)

We computed the robot's center of mass (CoM) during push-off as the mass-weighted average of the centers of mass of the 6 rigid components shown in Fig. S13:

$$\text{CoM} = \left( \frac{\sum_{i=1}^{n} m_i x_i}{\sum_{i=1}^{n} m_i}, \frac{\sum_{i=1}^{n} m_i y_i}{\sum_{i=1}^{n} m_i} \right), \tag{S1}$$

where $m_i$ is the mass of component $i$ and $(x_i, y_i)$ is the planar location of its center of mass. The component CoM locations were obtained from digitized joint coordinates (points $O$–$E$ in Fig. S13) as follows:

$$x_1 = \frac{x_O + x_A}{2}, \qquad y_1 = \frac{y_O + y_A}{2}, \tag{S2}$$

$$x_2 = \frac{x_A + x_B}{2}, \qquad y_2 = \frac{y_A + y_B}{2}, \tag{S3}$$

$$x_3 = x_B, \qquad y_3 = y_B, \tag{S4}$$

$$x_4 = \frac{x_C + x_D}{2}, \qquad y_4 = \frac{y_C + y_D}{2}, \tag{S5}$$

$$x_5 = \frac{x_D + x_E}{2}, \qquad y_5 = \frac{y_D + y_E}{2}, \tag{S6}$$

(S7)

For the SMA coil element, we defined the unit vector from $B$ to $E$ as

$$\mathbf{u}_{BE} = \left( \frac{x_E - x_B}{\|\mathbf{v}_{BE}\|}, \frac{y_E - y_B}{\|\mathbf{v}_{BE}\|} \right) \tag{S8}$$

and approximated the CoM of the spring–tendon segment as the midpoint between point $E$ and the point located a distance $l$ from $B$ along $\mathbf{u}_{BE}$:

$$x_6 = \frac{(x_B + l\,u_{BE,x}) + x_E}{2}, \qquad y_6 = \frac{(y_B + l\,u_{BE,y}) + y_E}{2}, \tag{S9}$$

where $l$ is the spring–tendon segment length defined in Fig. S13.

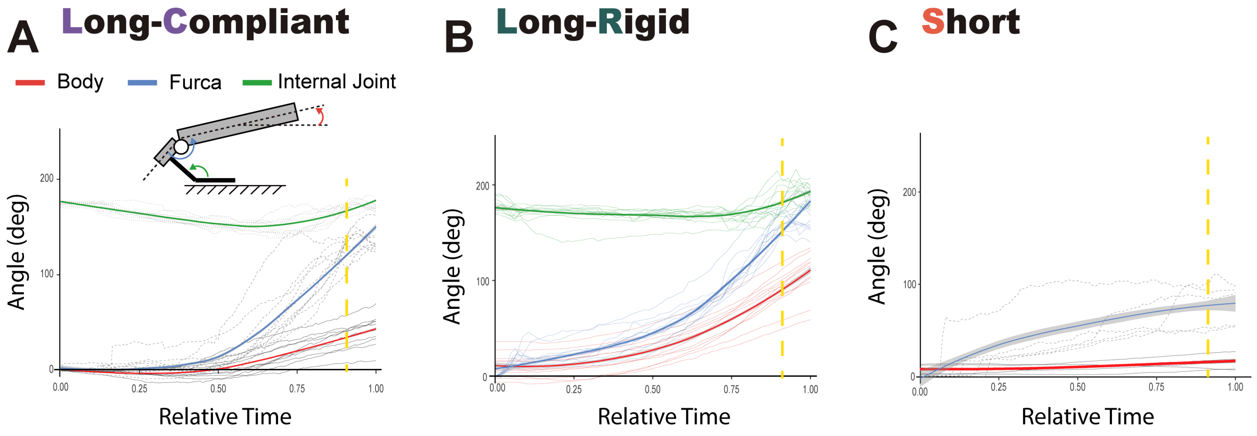


**Figure S1**: **Angular displacement of the body, furca, and internal furca joint.** (A) Long-compliant springtail. (B) Long-rigid springtail. (C) Short springtail.

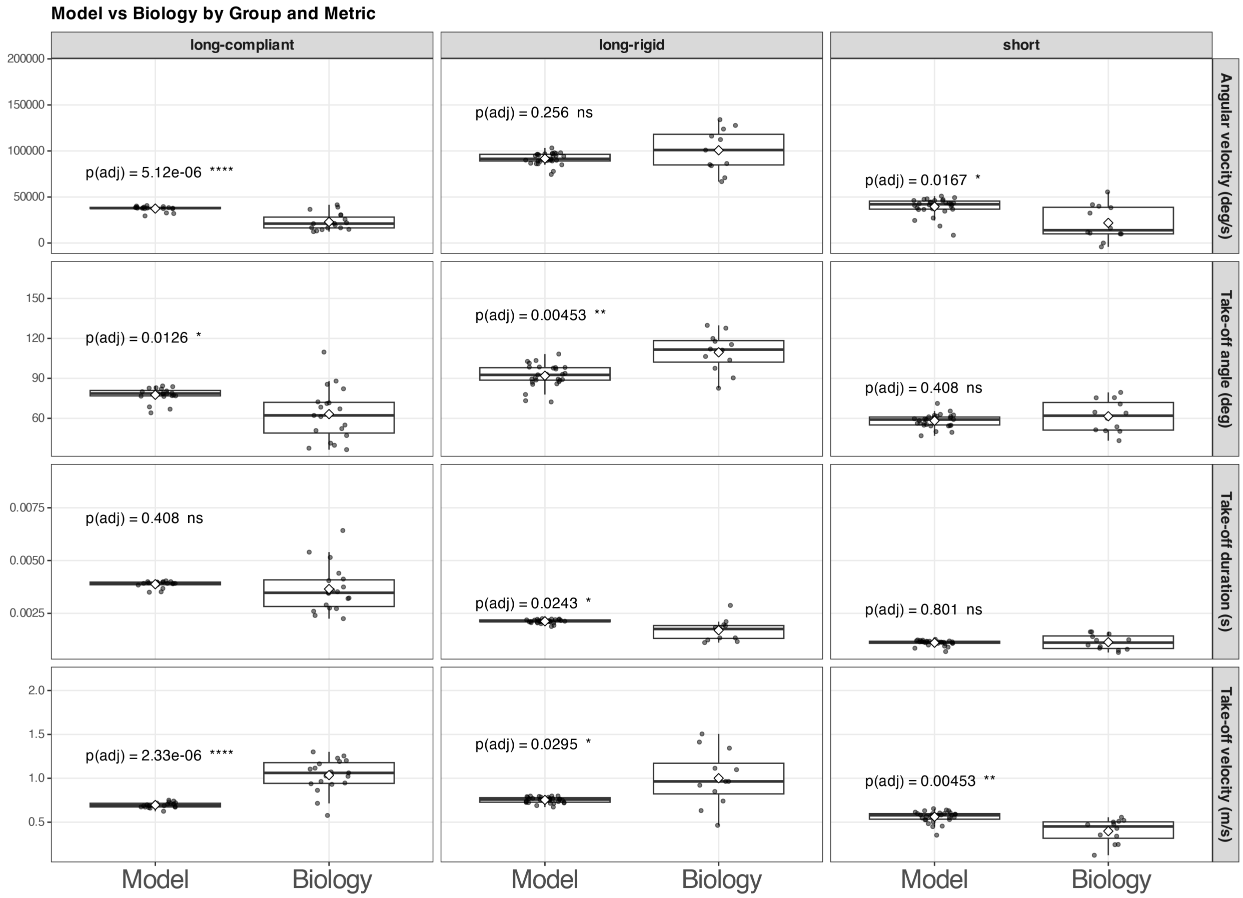


**Figure S2**: **Agreement between observed springtail kinematics and the mathematical model across morphotypes.** Columns correspond to morphotype (long-compliant, long-rigid, short). Rows report (top to bottom) body angular velocity at takeoff, takeoff angle, takeoff duration, and takeoff speed. For each panel, distributions of biological measurements and model outputs are shown as boxplots with individual points. Adjusted $p$-values for within-morphotype comparisons between biological data and model predictions are reported in each panel (test and correction described in Methods).

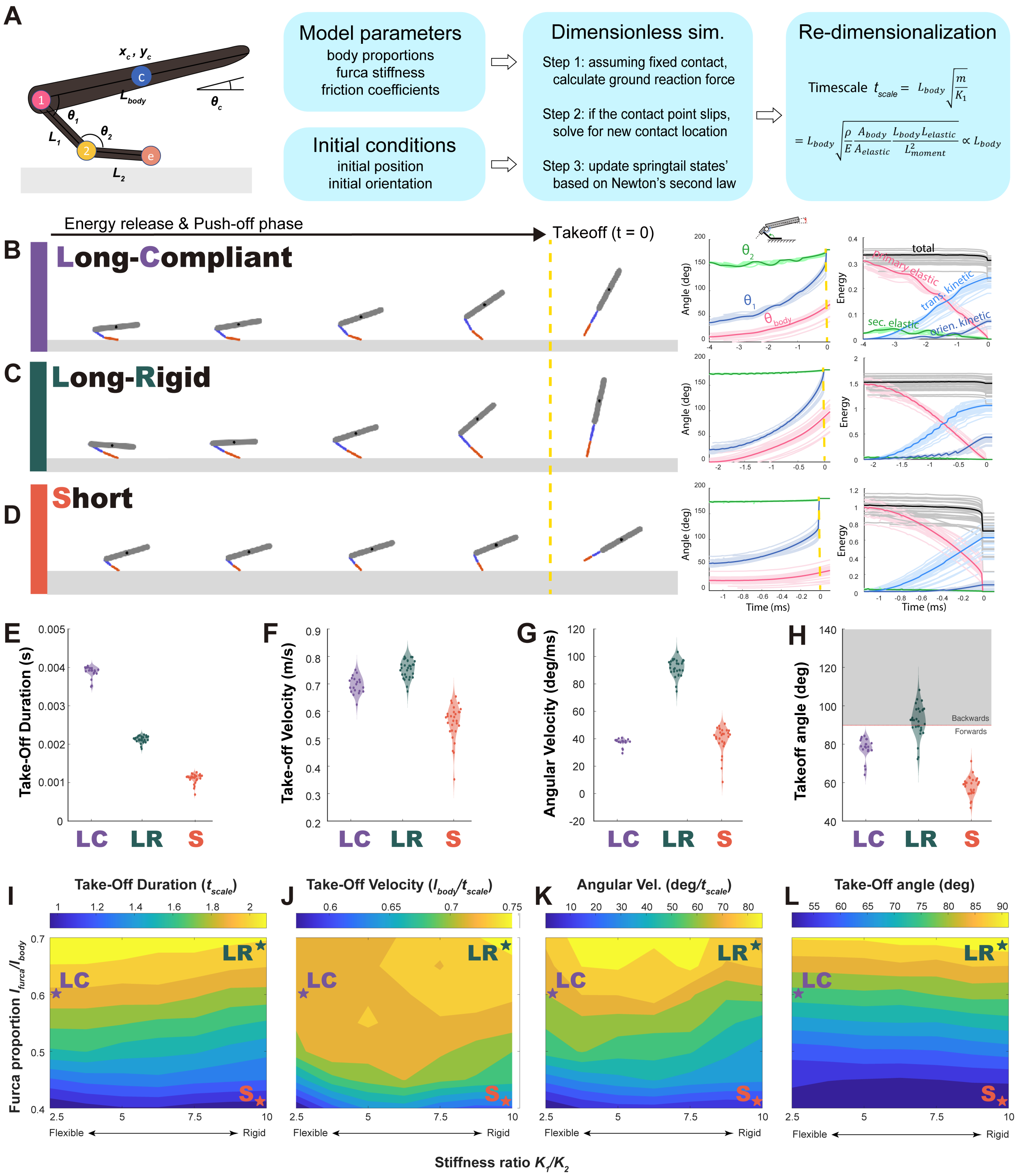

**Figure S3**: **Mathematical model captures the mechanism and performance of springtail jumping.** (A) Schematic of the planar rigid-body model with a two-joint furca represented by torsional springs. (B-D) Representative simulations for long-compliant, long-rigid, and short morphotypes, showing snapshots, joint-angle trajectories, and energy time series. Distributions of simulated contact duration, translational speed, angular velocity, and takeoff angle shown in dimensional form for comparison to Fig. 3. (I-L) Parameter sweeps showing how dimensionless jump metrics vary with furca length proportion and joint stiffness ratio.

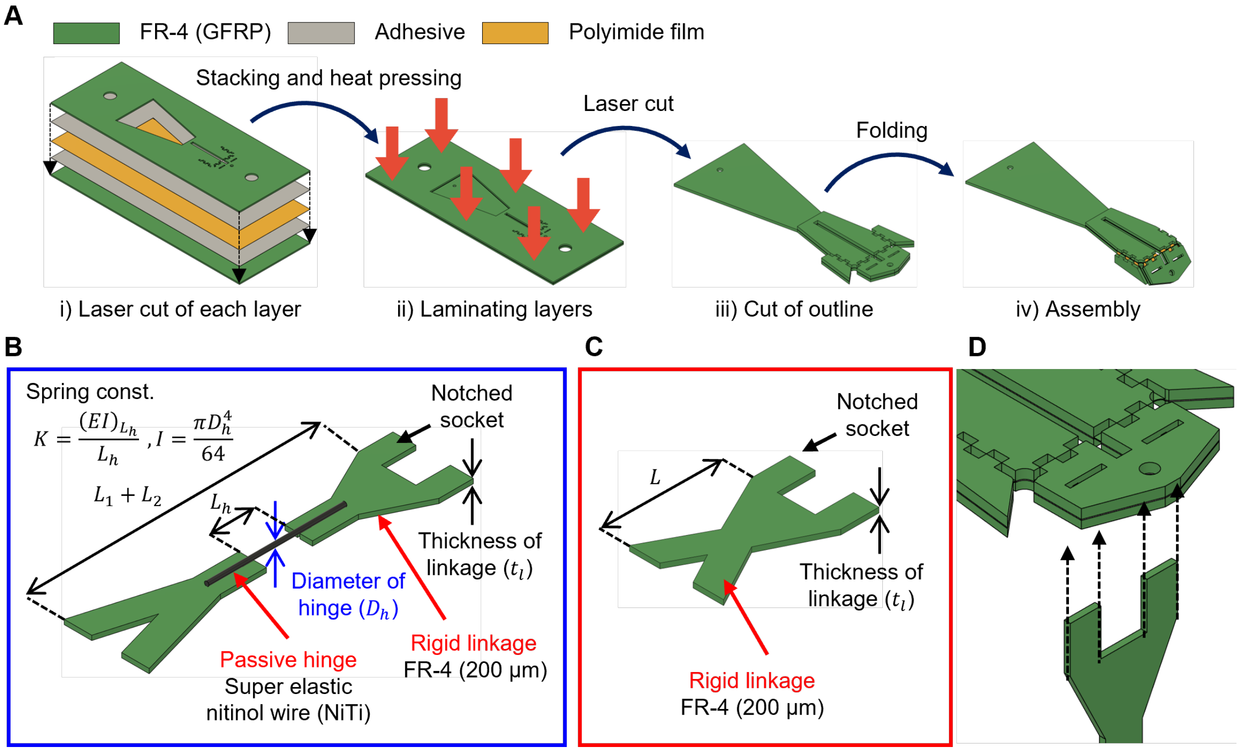


**Figure S4**: **Fabrication workflow for the jumping robot and robo-furca attachment.** (A) Folded composite body fabrication showing the layered laminate and hinge regions. (B) Long robo-furca with two-link geometry and hinge. (C) Short robo-furca fabricated as a single GFRP element. (D) Mechanical interface between the robot body and the robo-furca.

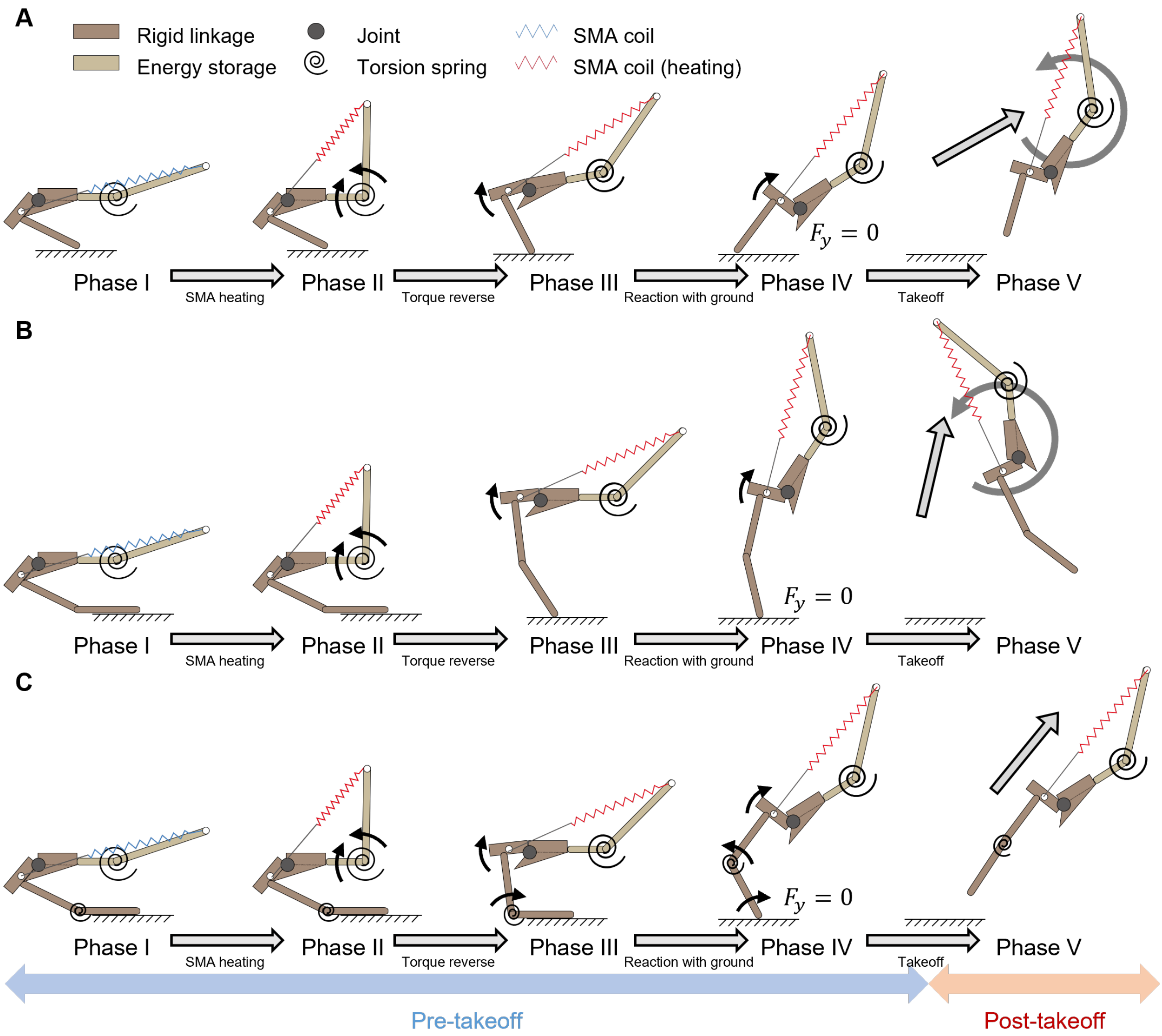


**Figure S5**: **Robo-springtail jumping sequence for three robo-furca morphotypes.** Image sequences illustrate the TRC-based actuation cycle for (A) short, (B) long-rigid, and (C) long-compliant robo-furcas. Panels show the progression from energy storage through rapid release and takeoff; symbols denote rigid links, compliant elements, and actuator state as indicated in the figure legend.

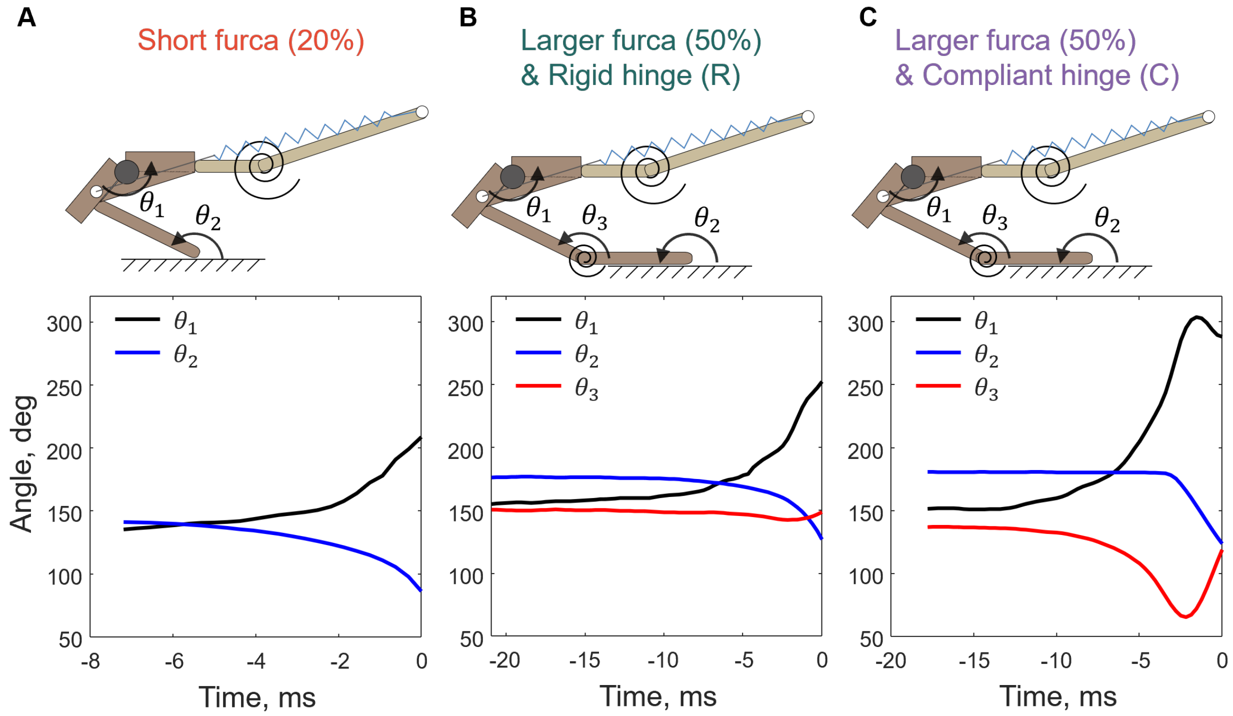


**Figure S6**: **Joint-angle trajectories during robo-springtail push-off for three robo-furca morphotypes.** (A) Short robo-furca (20% BL). (B) Long robo-furca (50% BL) with rigid hinge. (C) Long robo-furca (50% BL) with compliant hinge. Schematics define joint angles ($\theta_1$–$\theta_3$), and plots report their time evolution during the final milliseconds of push-off.

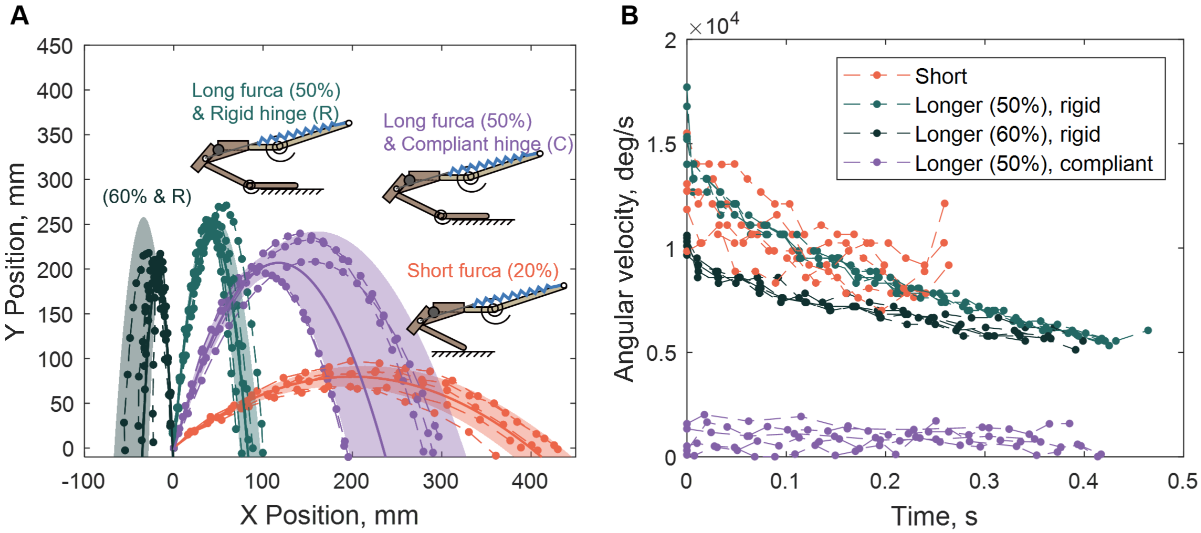


**Figure S7**: **Robo-springtail jump trajectories and airborne rotation depend on robo-furca morphology.** (A) Sagittal-plane trajectories for short, long-rigid, and long-compliant robo-furcas; markers show tracked positions over time. (B) Airborne angular velocity profiles following takeoff for the same conditions.

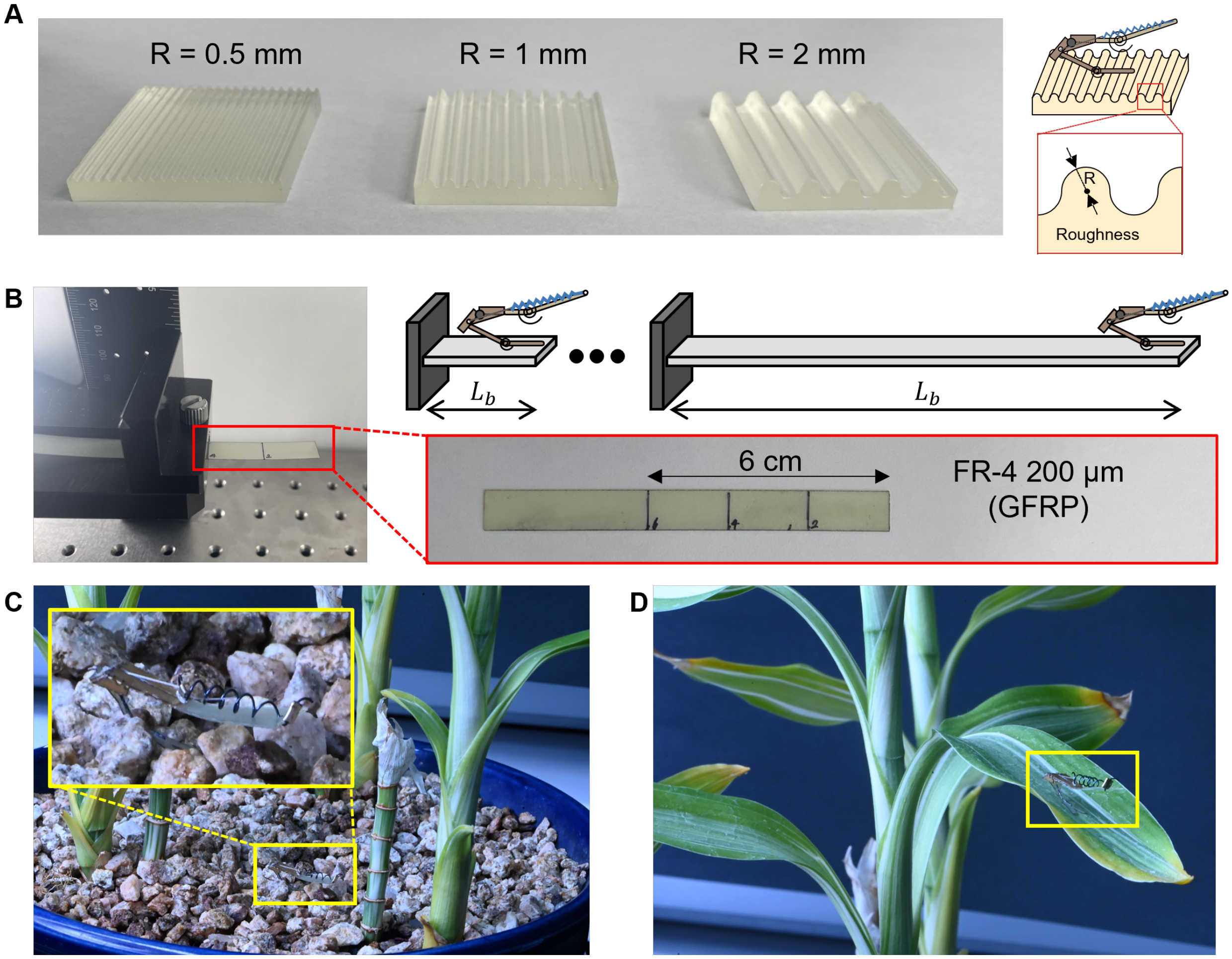


**Figure S8**: **Substrates used to test robo-springtail takeoff on uneven and compliant terrain.** (A) Quantified rough substrates fabricated with defined curvature radius $R$. (B) Quantified flexible substrates implemented as cantilever springboards with controlled beam length ($L_b$). (C) Natural rough substrate (gravel) used as an outdoor analog. (D) ) Natural compliant substrate (leaf) used as an outdoor analog.

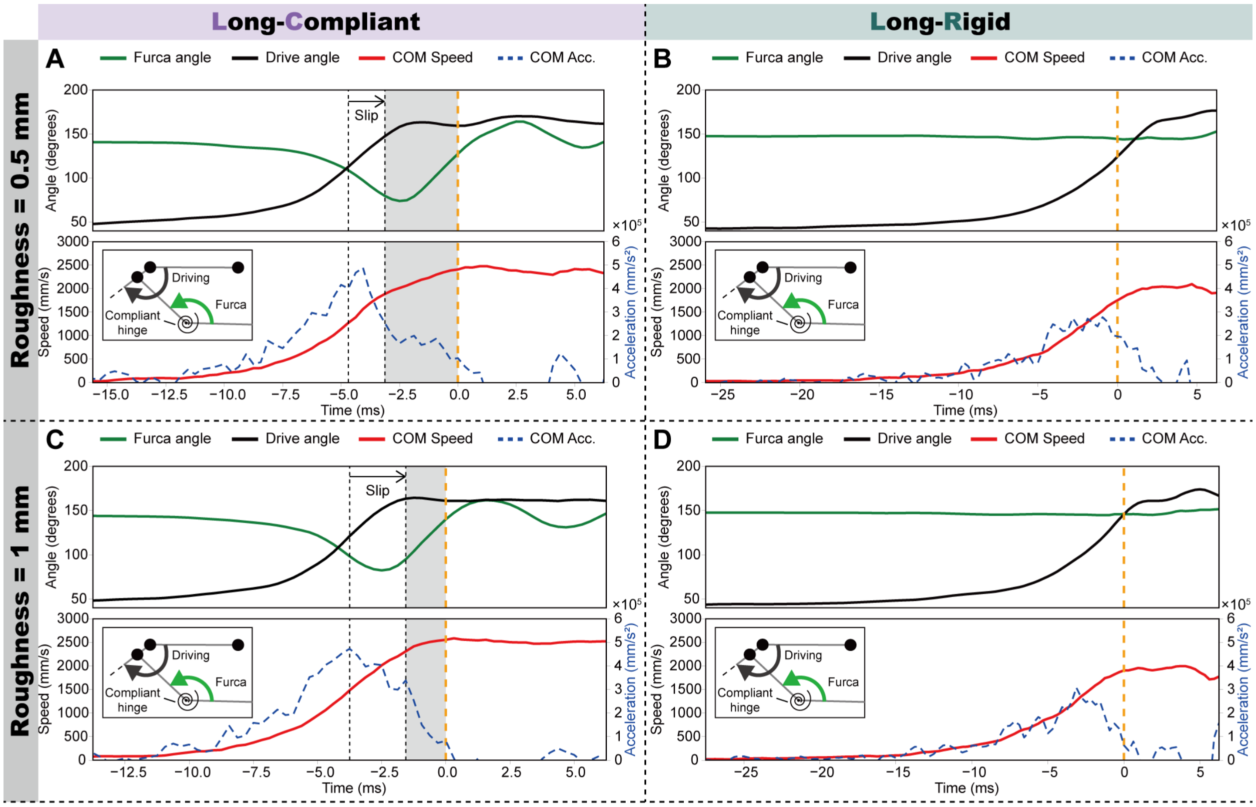


**Figure S9**: **Robo-springtail joint kinematics and body speed during push-off on 3D-printed uneven substrates.** Drive angle and internal robo-furca angle are plotted alongside the local substrate height at the contact point and the robot center-of-mass (CoM) speed and acceleration (line styles as indicated in the legend). (A, B) Roughness radius $R = 0.5$ mm for long-compliant and long-rigid robo-furcas. (C, D) $R = 1$ mm for long-compliant and long-rigid robo-furcas. The vertical line marks takeoff.

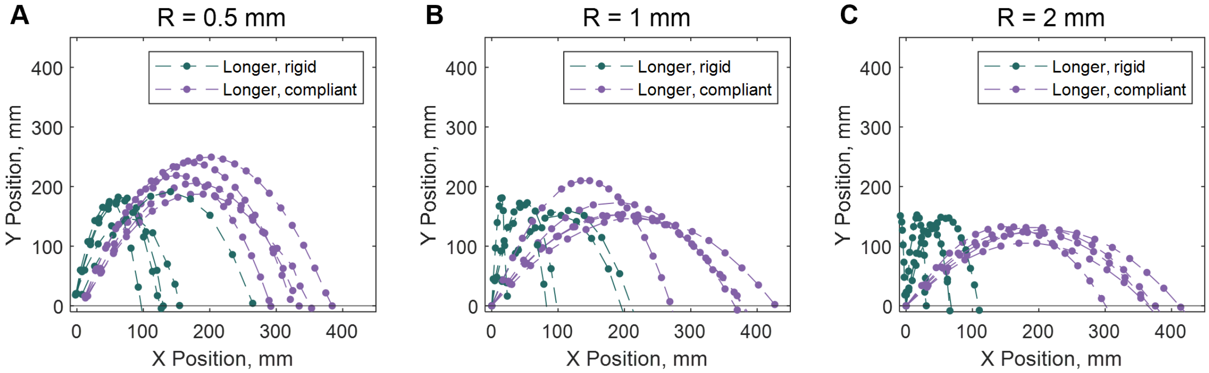


**Figure S10**: **Robo-springtail CoM trajectories on rough substrates of increasing curvature.** (A) $R = 0.5$ mm, (B) $R = 1$ mm, and (C) $R = 2$ mm.

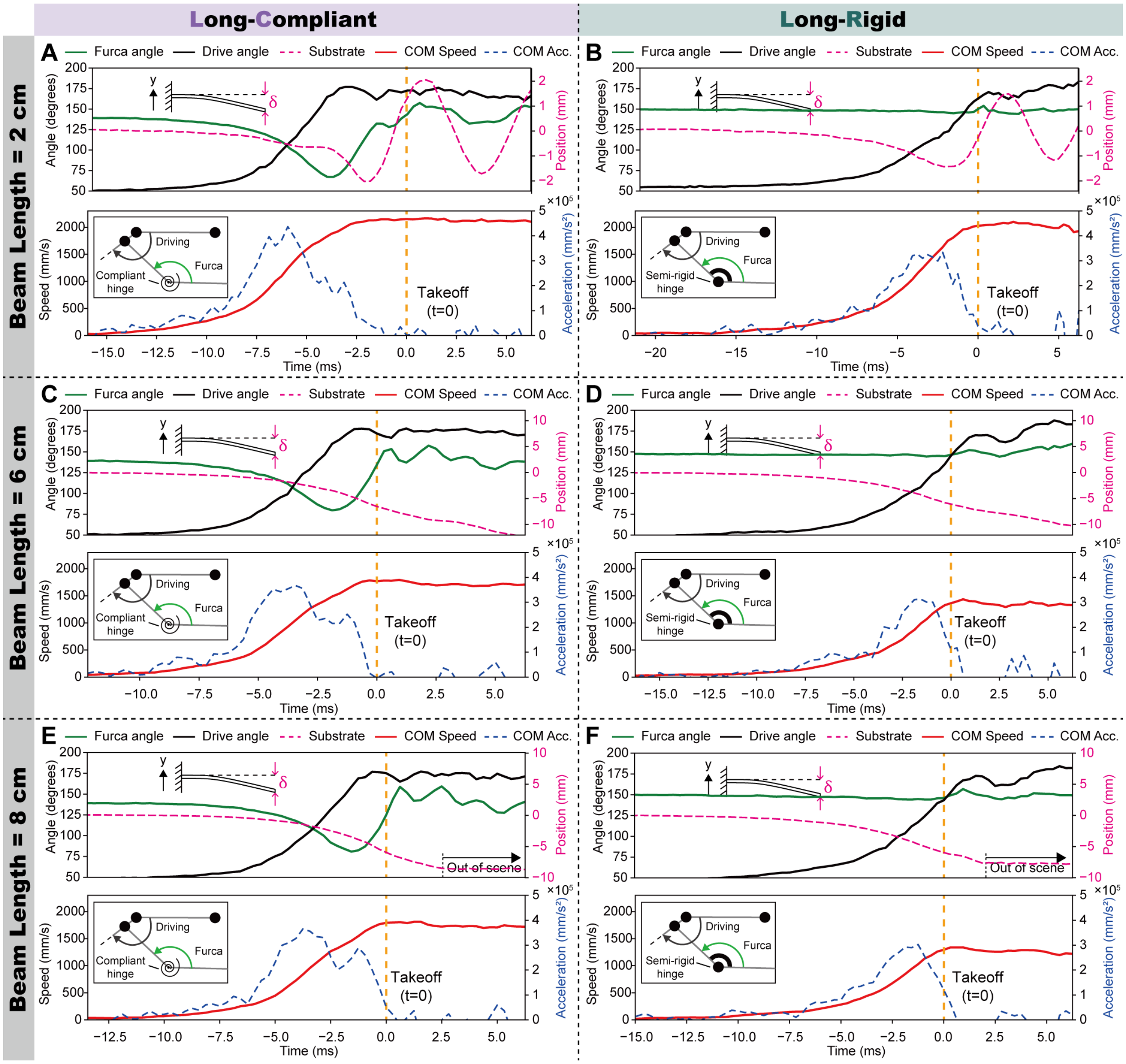


**Figure S11**: **Robo-springtail joint kinematics and body speed during push-off on cantilever springboards.** (A, B) Beam length = 2 cm. (C, D) Beam length = 6 cm. (E, F) Beam length = 8 cm. The vertical line marks takeoff.

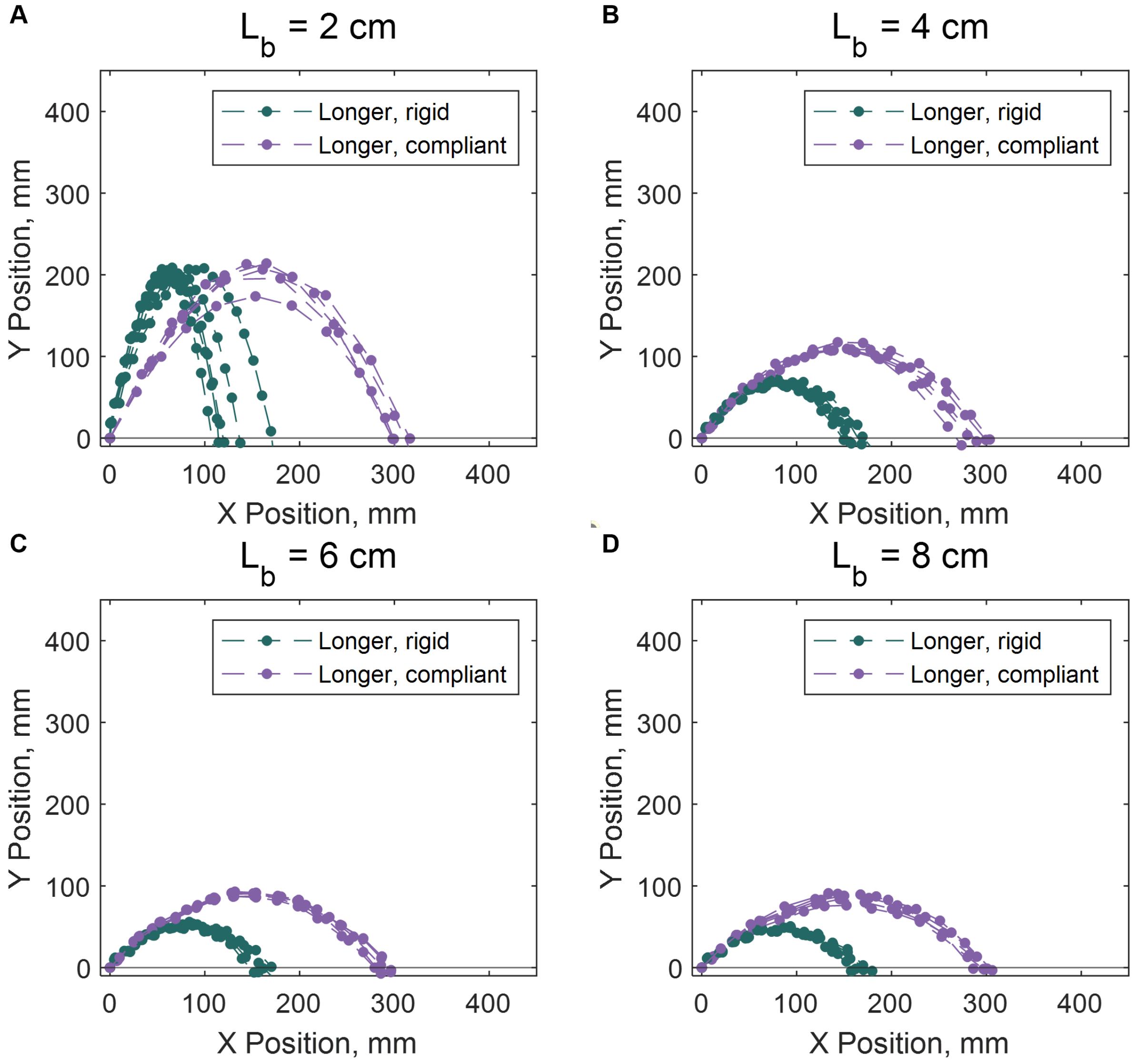


**Figure S12**: **Robo-springtail trajectories on springboards of increasing beam length.** (A) 2 cm, (B) 4 cm, (C) 6 cm, and (D) 8 cm beam lengths.

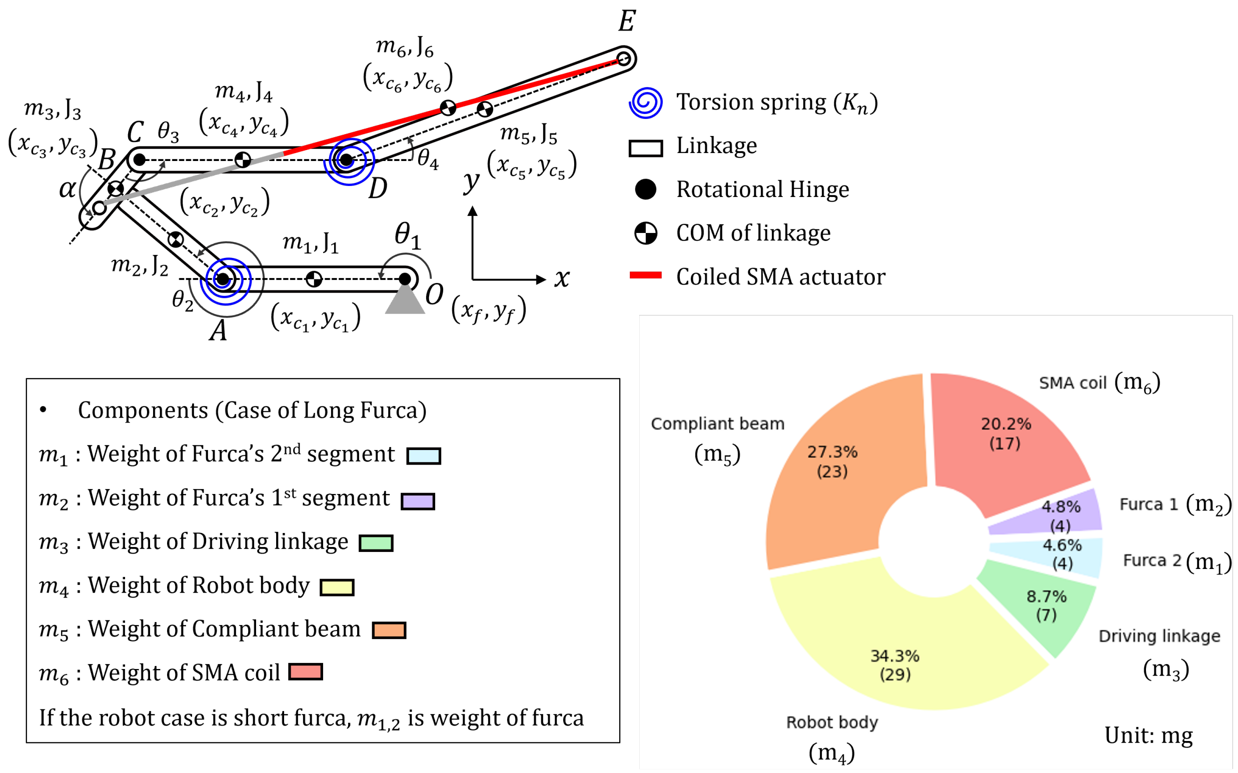


**Figure S13**: **Robo-springtail kinematic model and mass distribution used for center-of-mass calculations.** Schematic defines the tracked joint locations and the component CoM locations used in Eqs. S1–S9. The pie chart reports the mass partitioning across major components (body, drive linkage, robo-furca segments, compliant element, and actuator) as used in the CoM computation

**Table S1**: **Literature survey of jumping-robot capabilities relevant to terrain-adaptive locomotion.** Robots are grouped by body mass scale (< 1 g, 1–10 g, 10–100 g). “O” denotes a capability demonstrated experimentally in the cited work; “X” denotes not demonstrated or not reported. Terrain adaptability lists the specific terrain conditions tested when available.

| Scale | Name | Untethered Power | Aerial Stabilization | Landing or righting | Repeatable operation | Hybrid locomotion | Terrain adaptability |
|---|---|---|---|---|---|---|---|
| < 1g | 3.4-mm Flea-sized robot (*49*) | X | X | X | X | O | X |
| | Magnetically actuated gearbox (*50*) | X | X | X | O | X | X |
| | Soft jumping robot (*51*) | X | X | X | X | X | X |
| | This work (Long-compliant) | X | O | X | X | X | Rough and compliant |
| | This work (Long-rigid) | X | X | X | X | X | X |
| | This work (Short) | X | X | X | X | X | X |
| | Springtail-inspired robot (*11*) | X | O | O | X | X | X |
| | 216mg insect-sized jumping robot (*52*) | O | X | O | O | X | Slope |
| | Springtail-inspired microrobot (*53*) | O | X | X | X | O | X |
| 1-10g | Flea robot 1 (*54*) | X | X | X | X | X | X |
| | Soft combustion robot (*55*) | X | X | X | O | O | X |
| | Insect-scale jumping robots (*56*) | O | X | X | O | X | X |
| | Springtail-inspired jumping robot (*37*) | X | O | O | O | O | Different friction |
| | Flea robot 3 (*57*) | X | X | X | X | X | X |
| | Moobot (*58*) | O | X | X | X | X | X |
| | EPFL 7g (*59*) | O | O | X | X | X | X |
| | Tribot (*60*) | O | X | O | O | O | Gravel |
| 10-100g | MSU jumper (*61*) | O | X | O | O | O | O |
| | Salto-1P (*62*) | O | O | O | O | X | Moving plate |
| | JumpROACH (*63*) | O | X | O | O | O | X |
| | Salto (*16*) | O | O | X | O | O | X |
| | Forg-inspired jumping robot (*64*) | O | X | X | O | X | X |

**Table S2**: **Counts by springtail family in the comparative morphology image dataset.** Counts correspond to the filtered set of lateral-view images in which the furca was fully extended ($n = 552$).

| **Family** | **Number of Individuals** |
|---|---|
| Isotomidae | 127 |
| Neanuridae | 80 |
| Tomoceridae | 69 |
| Sminthuridae | 55 |
| Entomobryidae | 54 |
| Dicyrtomidae | 45 |
| Katiannidae | 30 |
| Hypogastruridae | 26 |
| Arrhopalitidae | 25 |
| Sminthurididae | 21 |
| Neelidae | 8 |
| Odontellidae | 7 |
| Oncopoduridae | 2 |
| Bourletiellidae | 2 |
| Brachystomellidae | 1 |

**Table S3**: Gaussian mixture model (GMM) fit to relative furca length distribution.

| **Parameter** | **Value** |
|---|---|
| Log-Likelihood | 332.451 |
| Number of Observations | 552 |
| Degrees of Freedom | 5 |
| BIC | 633.3 |
| ICL | 617.4 |

**Table S4**: **Scaling of relative furca length within the two mixture-model groups.** Linear regressions relate relative furca length (furca length/body length) to body length (mm), fit separately for individuals assigned to the small-furca (Peak 1) and long-furca (Peak 2) groups based on the mixture-model partition.

| Parameter | Peak 1 (Short Furcas) | Peak 2 (Long Furcas) |
|---|---|---|
| Intercept ($\beta_0$) | -0.027<br>$p = 0.334$ | 0.080<br>$p = 0.00572$ |
| Slope ($\beta_1$) | 0.181<br>$p < 2e - 16$ | 0.435<br>$p < 2e - 16$ |
| Residual Standard Error | 0.096 | 0.223 |
| Multiple R-squared | 0.739 | 0.753 |
| Adjusted R-squared | 0.737 | 0.752 |
| F-statistic | 441.8<br>$p < 2.2e - 16$ | 1193<br>$p < 2.2e - 16$ |
| Degrees of Freedom | 156 | 392 |

**Table S5**: **Top-view jump distance and direction across species and morphotypes.** Values are mean ± SD. *N* is the number of analyzed jumps. Jump length is reported in mm and normalized by body length (BL). Jump direction is the circular mean takeoff heading relative to initial body orientation (0° forward; 270° to 90° is a forward directed jump; Methods). Morphotypes: LC (long-compliant), LR (long-rigid), S (short).

| Species | Morphotype | *N* | Jump length (mm) | Jump length (BL) | Heading (°) |
|---|---|---|---|---|---|
| *Ceratophysella* spp. | S | 29 | 18.60 ± 5.01 | 15.88 ± 3.77 | 0.84 ± 5.50 |
| *F. candida* | S | 24 | 18.30 ± 5.28 | 12.37 ± 4.10 | 355.52 ± 25.83 |
| *D. minuta* | LR | 20 | 48.15 ± 24.85 | 28.16 ± 14.31 | 148.83 ± 90.20 |
| *H. sauteri* | LC | 22 | 41.60 ± 15.89 | 15.19 ± 5.79 | 334.47 ± 31.78 |
| *P. nigritus* | LC | 20 | 49.94 ± 16.05 | 15.79 ± 4.74 | 349.67 ± 50.53 |

**Table S6**: **Post hoc comparisons of relative jump distance across species.** Tukey's HSD results follow a one-way ANOVA on jump distance normalized by body length (BL).

| **Comparison** | **Mean Difference** | **Lower CI** | **Upper CI** | **p-value** |
|---|---|---|---|---|
| *D. minuta - Ceratophysella spp.* | 12.29 | 3.01 | 21.57 | 0.0037 |
| *F. candida - Ceratophysella spp.* | -3.50 | -13.43 | 6.42 | 0.8591 |
| *H. sauteri - Ceratophysella spp.* | -0.69 | -12.06 | 10.68 | 0.9998 |
| *P. nigritus - Ceratophysella spp.* | -0.09 | -9.37 | 9.19 | 1.0000 |
| *F. candida - D. minuta* | -15.79 | -24.14 | -7.44 | 0.0000 |
| *H. sauteri - D. minuta* | -12.98 | -23.00 | -2.95 | 0.0049 |
| *P. nigritus - D. minuta* | -12.38 | -19.96 | -4.80 | 0.0002 |
| *H. sauteri - F. candida* | 2.82 | -7.81 | 13.44 | 0.9455 |
| *P. nigritus - F. candida* | 3.41 | -4.94 | 11.77 | 0.7816 |
| *P. nigritus - H. sauteri* | 0.60 | -9.43 | 10.62 | 0.9998 |

**Table S7**: **Pairwise Watson–Williams tests comparing mean jump direction across species.** Tests compare circular means of top-view jump direction (degrees) using the Watson–Williams procedure; mean directions for each species are listed in the rightmost column.

| **Comparison** | **F-Statistic** | **Degrees of Freedom** | **p-value** | **Mean Direction (°)** |
|---|---|---|---|---|
| *Ceratophysella spp.* vs *D. minuta* | 37.22 | 1, 28 | 1.40e-06 | 0.84 (*Ceratophysella spp.*), 148.83 (*D. minuta*) |
| *Ceratophysella spp.* vs *F. candida* | 0.39 | 1, 22 | 0.54 | 0.84 (*Ceratophysella spp.*), 355.52 (*F. candida*) |
| *Ceratophysella spp.* vs *H. sauteri* | 6.14 | 1, 16 | 0.02 | 0.84 (*Ceratophysella spp.*), 334.47 (*H. sauteri*) |
| *Ceratophysella spp.* vs *P. nigritus* | 0.54 | 1, 28 | 0.47 | 0.84 (*Ceratophysella spp.*), 349.67 (*P. nigritus*) |
| *D. minuta* vs *F. candida* | 38.42 | 1, 32 | 6.12e-07 | 148.83 (*D. minuta*), 355.52 (*F. candida*) |
| *D. minuta* vs *H. sauteri* | 100.05 | 1, 26 | 2.11e-10 | 148.83 (*D. minuta*), 334.47 (*H. sauteri*) |
| *D. minuta* vs *P. nigritus* | 37.54 | 1, 38 | 3.81e-07 | 148.83 (*D. minuta*), 349.67 (*P. nigritus*) |
| *F. candida* vs *H. sauteri* | 2.60 | 1, 20 | 0.12 | 355.52 (*F. candida*), 334.47 (*H. sauteri*) |
| *F. candida* vs *P. nigritus* | 0.16 | 1, 32 | 0.69 | 355.52 (*F. candida*), 349.67 (*P. nigritus*) |
| *H. sauteri* vs *P. nigritus* | 0.65 | 1, 26 | 0.43 | 334.47 (*H. sauteri*), 349.67 (*P. nigritus*) |

**Table S8**: **Lateral-view takeoff kinematics by species and morphotype.** Values are mean ± s.d.; $N$ is the number of jumps. Takeoff duration is the interval from furca release to loss of ground contact. Body angular velocity is computed during takeoff.

| **Species** | **Morphotype** | $N$ | **Duration (ms)** | **Speed (m/s)** | **Body Ang. Vel. (°/ms)** | **Angle (°)** |
|---|---|---|---|---|---|---|
| *Ceratophysella spp.* | short | 5 | 0.81 ± 0.13 | 0.26 ± 0.09 | 7.64 ± 6.61 | 54.2 ± 8.1 |
| *F. candida* | short | 11 | 1.37 ± 0.25 | 0.50 ± 0.04 | 32.02 ± 18.37 | 66.8 ± 11.8 |
| *D. minuta* | long-rigid | 12 | 1.70 ± 0.50 | 1.00 ± 0.31 | 100.77 ± 22.40 | 109.5 ± 14.3 |
| *P. nigritus* | long-compliant | 10 | 3.39 ± 1.23 | 0.99 ± 0.21 | 20.46 ± 8.40 | 53.5 ± 17.1 |
| *H. sauteri* | long-compliant | 14 | 3.95 ± 0.87 | 1.09 ± 0.14 | 25.65 ± 8.94 | 73.9 ± 16.9 |

**Table S9**: **Statistical tests comparing kinematic variables across morphotypes.** We report one-way ANOVA ($F$) or Kruskal–Wallis ($\chi^2$) test statistics. For ANOVA, SS and MS denote sums of squares and mean squares.

| **Metric** | **Test** | **Df** | **SS** | **MS** | **Statistic** | **p-value** |
|---|---|---|---|---|---|---|
| Takeoff angle | ANOVA | 2 | 19135 | 9567 | $F = 35.78$ | $1.23 \times 10^{-9}$ |
| Takeoff velocity | ANOVA | 2 | 3383846 | 1691923 | $F = 35.51$ | $1.36 \times 10^{-9}$ |
| Jump duration | Kruskal–Wallis | 2 | – | – | $\chi^2 = 32.34$ | $9.50 \times 10^{-8}$ |
| Body angular velocity | Kruskal–Wallis | 2 | – | – | $\chi^2 = 25.74$ | $2.57 \times 10^{-6}$ |

**Table S10**: **Pairwise comparisons across morphotypes.**

| **Parameter** | **Comparison** | **p-value** |
|---|---|---|
| TakeOffAngle | long-compliant vs long-rigid | 1.9e-06 ** |
| | long-compliant vs short | 1.0 |
| | short vs long-rigid | 2.2e-06 ** |
| TakeOffVelocity | long-compliant vs long-rigid | 1.0 |
| | long-compliant vs short | 4.3e-08 ** |
| | short vs long-rigid | 6.7e-05 ** |
| JumpDuration | long-compliant vs long-rigid | 3.2e-05 ** |
| | long-compliant vs short | 1.2e-05 ** |
| | short vs long-rigid | 0.022 * |
| BodyAngleVelocity | long-compliant vs long-rigid | 4.3e-08 ** |
| | long-compliant vs short | 1.0 |
| | short vs long-rigid | 2.2e-06 ** |

**Table S11**: **Comparison of linear mixed-effects and simple linear models.** The mixed model includes species as a random effect, yielding more conservative intercept estimates. Despite fewer residual degrees of freedom, the mixed model provides a better fit.

| Model | Parameter | Estimate | Std. Error | $p$-value |
|---|---|---|---|---|
| **Fixed effects** | | | | |
| Mixed (LMER) | Intercept | −28.25 | 16.14 | 0.10 |
| | MaxIntFurcaAngle | 0.88 | 0.12 | $2.7 \times 10^{-7}$ |
| Simple (LM) | Intercept | −55.98 | 11.48 | $3.6 \times 10^{-5}$ |
| | MaxIntFurcaAngle | 1.11 | 0.09 | $6.7 \times 10^{-13}$ |
| **Random effects (mixed model only)** | | | | |
| Species (Intercept) | | 80.89 | 8.99 | |
| Residual | | 116.21 | 10.78 | |
| **Model fit statistics** | | | | |
| Mixed model: REML = 236.7, ICC = 0.41, $N = 31$ (3 species) | | | | |
| Simple model: $R^2 = 0.836$, adj. $R^2 = 0.830$, residual SE = 11.89, $F(1, 29) = 147.6$ | | | | |
| Residuals: Shapiro–Wilk test $p > 0.25$ | | | | |

**Table S12**: **Jumping robot specifications and performance metrics across scales (literature survey).** Metrics are reported as in the cited sources; dashes indicate values not reported.

| Name | Mass (g) | Length (mm) | Leg length (mm) | Jump height (m) | Jump distance (m) | Speed (m/s) | Push-off duration (ms) | Energy (mJ) |
|---|---|---|---|---|---|---|---|---|
| 3.4-mm Flea-sized robot (*49*) | 0.012 | 3.4 | 0.3 | 0.104 | 0.296 | 1.74 | 10 | 0.018 |
| Magnetically actuated gearbox (*50*) | 0.025 | 3.1 | 3.1 | 0.119 | 0.218 | 2.3 | 0.75 | 0.067 |
| Soft jumping robot (*51*) | 0.08 | 56 | 28 | 0.062 | 0.041 | 1.12 | 21 | 0.050 |
| This work (Long-compliant) | 0.08 | 20 | 10 | 0.216 | 0.250 | 2.135 | 16.054 | 0.182 |
| This work (Long-rigid) | 0.08 | 20 | 10 | 0.256 | 0.083 | 2.741 | 19.054 | 0.301 |
| This work (Short) | 0.08 | 20 | 4 | 0.081 | 0.402 | 2.228 | 7.946 | 0.198 |
| Springtail-inspired robot (*11*) | 0.1 | 20 | 11 | 0.46 | 0.335 | 3 | - | 0.45 |
| 216mg insect-sized jumping robot (*52*) | 0.216 | 24 | 12 | 0.24 | - | 2.1 | - | 0.476 |
| Springtail-inspired microrobot (*53*) | 0.98 | 21 | 10 | - | - | 3.171 | 7 | 4.927 |
| Flea robot 1 (*54*) | 1.104 | 20 | 31 | 0.64 | - | 4.2 | 8 | 9.737 |
| Soft combustion insect-scale robot (*55*) | 1.6 | 29 | 7 | 0.59 | 0.16 | 2.5 | 5 | 0.24 |
| Insect-scale jumping robots (*56*) | 1.68 | 23.1 | 3.5 | 0.89 | - | 4.2 | 3.5 | 14.818 |
| Springtail-inspired jumping robot (*37*) | 2.2 | 61 | 22.3 | 0.62 | 1.4 | 4.21 | 14.15 | 19.497 |
| Flea robot 3 (*57*) | 2.25 | 30 | 60 | 1.2 | - | 7 | 8 | 55.125 |
| Moobot (*58*) | 6 | 50 | 44 | 0.63 | - | 3.5 | - | 36.75 |
| EPFL 7g (*59*) | 7 | 50 | 100 | 1.38 | 0.79 | 5.9 | 19 | 121.835 |
| Tribot (*60*) | 9.7 | 58 | 44 | 0.14 | 0.23 | 1.65 | - | 13.204 |
| MSU jumper (*61*) | 20 | 65 | 40 | 0.55 | - | 3.34 | - | 111.556 |
| Salto-1P (*62*) | 98.1 | 150 | 144 | 1.25 | 2 | 4.95 | 57 | 1201.848 |
| JumpROACH (*63*) | 99 | 120 | 95 | 1.5 | 0.8 | 5.39 | 15 | 1438.079 |
| Salto (*16*) | 100 | 150 | 150 | 1.008 | - | 4.44 | 60 | 985.68 |
| Frog-inspired jumping robot (*64*) | 100.7 | 100 | 160 | 1.3 | 0.3 | 4.9 | 30 | 1208.904 |

**Table S13**: **Takeoff speed of biological and robotic jumpers on compliant substrates.** Substrate stiffness is the effective vertical stiffness at the contact point (N/m) as reported or inferred in the cited study; "Rigid" denotes a noncompliant reference surface.

| Jumper | Mass (g) | Substrate Stiffness (N/m) | Takeoff Velocity (m/s) | Note |
|---|---|---|---|---|
| *Robots* | | | | |
| 4g LaMSA Jumper (R=0) | 4.43 | 4963 | 2.63[a] | Substrate mass: 0.02216 kg |
| 4g LaMSA Jumper (R=2) | 4.43 | 4963 | 2.42[a] | Substrate mass: 0.02216 kg |
| 4g LaMSA Jumper (R=0) | 4.43 | 4963 | 2.36[a] | Substrate mass: 0.00295 kg |
| 4g LaMSA Jumper (R=2) | 4.43 | 4963 | 2.37[a] | Substrate mass: 0.00295 kg |
| 4g LaMSA Jumper (R=0) | 4.43 | 4963 | 2.55[a] | Substrate mass: 0.00148 kg |
| 4g LaMSA Jumper (R=2) | 4.43 | 4963 | 2.45[a] | Substrate mass: 0.00148 kg |
| 4g LaMSA Jumper (R=0) | 4.43 | 1489 | 2.52[a] | Substrate mass: 0.02216 kg |
| 4g LaMSA Jumper (R=4) | 4.43 | 1489 | 2.11[a] | Substrate mass: 0.02216 kg |
| 4g LaMSA Jumper (R=0) | 4.43 | 1489 | 1.70[a] | Substrate mass: 0.00295 kg |
| 4g LaMSA Jumper (R=4) | 4.43 | 1489 | 1.65[a] | Substrate mass: 0.00295 kg |
| 4g LaMSA Jumper (R=0) | 4.43 | 1489 | 1.61[a] | Substrate mass: 0.00148 kg |
| 4g LaMSA Jumper (R=4) | 4.43 | 1489 | 1.80[a] | Substrate mass: 0.00148 kg |
| 4g LaMSA Jumper (R=0) | 4.43 | 744.5 | 2.62[a] | Substrate mass: 0.02216 kg |
| 4g LaMSA Jumper (R=2) | 4.43 | 744.5 | 2.40[a] | Substrate mass: 0.02216 kg |
| 4g LaMSA Jumper (R=0) | 4.43 | 744.5 | 1.91[a] | Substrate mass: 0.00295 kg |
| 4g LaMSA Jumper (R=2) | 4.43 | 744.5 | 1.79[a] | Substrate mass: 0.00295 kg |
| 4g LaMSA Jumper (R=0) | 4.43 | 744.5 | 1.48[a] | Substrate mass: 0.00148 kg |
| 4g LaMSA Jumper (R=2) | 4.43 | 744.5 | 1.45[a] | Substrate mass: 0.00148 kg |
| Salto Robot | 108 | 1000 | 2.00[b] | Damping: $\zeta \approx 0.1$ |
| Salto Robot | 108 | 1000 | 1.97[b] | Damping: $\zeta \approx 1.0$ |
| Salto Robot | 108 | 200 | 1.91[b] | Damping: $\zeta \approx 0.1$ |
| Salto Robot | 108 | 200 | 1.74[b] | Damping: $\zeta \approx 1.0$ |
| Salto Robot | 108 | 100 | 1.80[b] | Damping: $\zeta \approx 0.1$ |
| Salto Robot | 108 | 100 | 1.60[b] | Damping: $\zeta \approx 1.0$ |
| Salto Robot | 108 | 85 | 1.74[b] | Damping: $\zeta \approx 0.1$ |
| Salto Robot | 108 | 85 | 1.62[b] | Damping: $\zeta \approx 1.0$ |

Cont'd

| Jumper | Mass (g) | Substrate Stiffness (N/m) | Takeoff Velocity (m/s) | Note |
|---|---|---|---|---|
| *Organisms* | | | | |
| White-cheeked Gibbon | 7700 | Rigid | 2.71[c] | Orthograde leaps |
| White-cheeked Gibbon | 7700 | 30000 | 1.88[c] | Orthograde leaps |
| White-cheeked Gibbon | 7700 | Rigid | 2.87[c] | Pronograde leaps |
| White-cheeked Gibbon | 7700 | 30000 | 2.69[c] | Pronograde leaps |
| Cuban Tree Frog | 22 | Rigid | 2.29[d] | - |
| Cuban Tree Frog | 22 | 95.24 | 2.02[d] | - |
| Cuban Tree Frog | 22 | 56.5 | 1.80[d] | - |
| Cuban Tree Frog | 22 | 43.29 | 1.47[d] | - |
| Green Anole Lizard | 2.05 | Rigid | 1.26[e] | - |
| Green Anole Lizard | 2.05 | 3.7 | 1.17[e] | - |
| Green Anole Lizard | 2.05 | 1.56 | 1.23[e] | - |
| Green Anole Lizard | 5.43 | Rigid | 1.37[e] | - |
| Green Anole Lizard | 5.43 | 3.7 | 1.25[e] | - |
| Green Anole Lizard | 5.43 | 1.56 | 1.09[e] | - |
| Fox squirrel | 829 | Rigid | 2.17[f] | - |
| Fox squirrel | 829 | 33.50 | 2.12[f] | - |
| Tree frog | 71.9 | 634 | 2.97[g] | - |
| Tree frog | 71.9 | 258 | 2.64[g] | - |
| Tree frog | 71.9 | 87 | 2.45[g] | - |
| Tree frog | 71.9 | 32 | 1.91[g] | - |

[a]Calculated from energy profile in Figure S11-S19 (*19*).

[b]Estimated from the plot in Fig. 6F (*23*).

[c]Calculated from kinetic energy and body weight in Table 1 (*24*).

[d]Estimated from the plot in Fig. 2B (*26*).

[e]Estimated from the plot in Fig. 1C and 1D (*25*).

[f]Data from https://zenodo.org/records/5140578 (*65*)

[g]Estimated from the plot in Figure 5C (*66*).

**Table S14**: **Hartigan's dip test rejects unimodality in relative furca length.** Dip test statistic and $p$-value are reported for the distribution of furca length normalized by body length ($n = 552$).

| **Test Statistic (D)** | **P-value** | **Conclusion** |
|---|---|---|
| 0.057443 | $p < 2.2e - 16$ | Not Unimodal |

**Table S15**: **Effective bending stiffness of cantilever springboards used as compliant substrates.** Bending stiffness (N/m) is the effective tip stiffness of the GFRP springboard used in the robot experiments for each beam length.

| **Beam length (cm)** | **Bending stiffness (N/m)** |
|---|---|
| 2 | 26.34 |
| 4 | 3.292 |
| 6 | 0.9754 |
| 8 | 0.4115 |

**Caption for Movie S1. Springtail jump kinematics and corresponding physics-based mathematical model output.**

[00:05] Springtails (Collembola) that use a spring-actuated mechanism to jump dozens of times their body length in the fraction of a second.

[00:25] A specialized structure called a furca for springtails to jump.

[00:36] Diverse body shape and furca morphology of springtails.

[00:48] Three groups of springtails that differed in furca size and flexibility: Short, Long-Rigid, and Long-Compliant.

[01:07] Kinematic movement and angle profiles of the springtail with short furca.

[01:20] Kinematic movement and angle profiles of the springtail with long-rigid furca.

[01:25] Kinematic movement and angle profiles of the springtail with long-compliant furca.

[01:41] Short, Long-Rigid, and Long-Compliant jumping mathematical model to support biological findings.

[01:46] Kinematic movement and angle profiles of the short furca model.

[01:55] Kinematic movement and angle profiles of the long-rigid furca model.

[02:03] Kinematic movement and angle profiles of the long-compliant furca model.

**Caption for Movie S2. Robo-springtail kinematics during push-off and full jump trajectories for three robo-furca designs.**

[00:05] Springtail-inspired robot equipped with various types of jumping appendage: Short, Long-Rigid, and Long-Compliant.

[00:09] Kinematic movement and angle profiles of the robot with short robo-furca.

[00:14] Kinematic movement and angle profiles of the robot with long-rigid robo-furca.

[00:20] Kinematic movement and angle profiles of the robot with long-compliant robo-furca.

[00:29] Full jumping trajectory of the robot with short robo-furca.

[00:40] Full jumping trajectory of the robot with long-rigid robo-furca.

[00:54] Full jumping trajectory of the robot with long-compliant robo-furca.

**Caption for Movie S3. Jumping of robots on the rough substrates.**

[00:05] Experimental setup.

[00:13] Comparison of the robot's kinematic movement between rigid and compliant robo-furca on rough substrate.

[00:54] Jumping of the robot with long-rigid robo-furca (R = 0.5 mm).

[01:03] Jumping of the robot with long-compliant robo-furca (R = 0.5 mm).

[01:14] Jumping of the robot with long-rigid robo-furca (R = 1 mm).

[01:23] Jumping of the robot with long-compliant robo-furca (R = 1 mm).

[01:34] Jumping of the robot with long-rigid robo-furca (R = 2 mm).

[01:43] Jumping of the robot with long-compliant robo-furca (R = 2 mm).

**Caption for Movie S4. Jumping of robots on the natural terrain (gravel).**

[00:05] Jumping of the robot with long-rigid robo-furca on gravel.

[00:12] Jumping of the robot with long-compliant robo-furca on gravel.

**Caption for Movie S5. Jumping of robots on the compliant substrates.**

[00:05] Experimental setup.

[00:13] Comparison of the robot's kinematic movement between rigid and compliant robo-furca on a compliant substrate.

[01:02] Jumping of the robot with long-rigid robo-furca (L = 2 cm).

[01:12] Jumping of the robot with long-compliant robo-furca (L = 2 cm).

[01:23] Jumping of the robot with long-rigid robo-furca (L = 4 cm).

[01:32] Jumping of the robot with long-compliant robo-furca (L = 4 cm).

[01:42] Jumping of the robot with long-rigid robo-furca (L = 6 cm).

[01:51] Jumping of the robot with long-compliant robo-furca (L = 6 cm).

[02:01] Jumping of the robot with long-rigid robo-furca (L = 8 cm).

[02:12] Jumping of the robot with long-compliant robo-furca (L = 8 cm).

**Caption for Movie S6. Jumping of robots on the natural terrain (leaf and pine needles).**

[00:10] Jumping of the robot with long-rigid robo-furca on a leaf.

[00:19] Jumping of the robot with long-compliant robo-furca on a leaf.

[00:30] Jumping of the robot with long-rigid robo-furca on pine needles.

[00:41] Jumping of the robot with long-compliant robo-furca on pine needles.
